# Emergent metabolic cooperation in communities of microbes that maximise growth rate by optimal protein expression

**DOI:** 10.64898/2026.10.28.762437

**Authors:** Maarten J. Droste, Robert Planqué, Frank J. Bruggeman

## Abstract

Many microbial communities involve metabolic exchanges between their members. We present a mathematical modelling formalism for studying the adaptation of metabolic strategies by microbes to external metabolites produced by themselves and their community partners. Each microbial species is assumed to maximise its growth rate by optimal allocation of finite biosynthetic resources for protein synthesis in response to their current environment. Each species therefore acts as a selfish optimiser. Because the environment is shaped by all species, their strategies are interdependent. The optimal state of the community is the set of metabolic strategies of its members that outcompetes (outgrows) all other possible sets of metabolic strategies—including the ones in which some species may not be present at all. Our mathematical formalism compacts the coupled optimisations by the single species into one optimisation problem that also infers the optimal community composition. We show with simple examples how it can be used to study the emergence of positive species-species interactions such as commensalism, mutualism, syntrophy, and division of labour. The formalism is a direct generalisation of the well-known resource allocation models for single species, but includes reaction kinetics; we show that observed positive ecological interactions emerge from ecological feedback, in which producers are inhibited by the byproducts they excrete. This paper illustrates that (metabolism-dominated) microbial communities may be viewed as one coupled system in which the optimal pathway choice of each member depends on that of all others. It is directly generalisable to multiple species, and should pave the way for studies of larger communities.

## 1 Introduction

Microbial communities contribute to chemical-element cycling on our planet [1], are vital for higher eukaryotes [2], and ferment many of our foods and beverages [3]. Positive and negative interactions between microbial species shape the composition of the community and determine its ecological services. One example is the anaerobic digestion of plant-derived lignocellulose in soils [4]. A key interaction in this community is that a catabolic byproduct of one microbe, e.g., the acetate made during sugar metabolism, acts as the nutrient source for the next microbe. Together, these microbes create a metabolic network, with individual cells acting as catalysts that compete and cooperate for metabolites through cross-feeding [4–6]. This paper focuses on such metabolism-dominated microbial communities [7].

Which metabolic product a microbe excretes results from the available nutrients and the expressed metabolic strategy. *Escherichia coli* can, for instance, aerobically respire glucose into carbon dioxide and water or convert it (partially) into acetate, while anaerobically it converts glucose via mixed acid fermentation into acetate, ethanol and formate. Whether acetate is formed under aerobic conditions has been shown to depend on the growth rate [8]. A critical growth rate exists beyond which acetate production commences, indicating a shift from respiration to acetate formation. This has been attributed to a fundamental limit on the allocation of finite biosynthetic resources for protein synthesis [9]. A higher growth rate demands a larger anabolic protein pool, leaving less biosynthetic resources for catabolism. Above a critical growth rate a microbe can only increase its growth rate if it has a catabolic strategy that can supply ATP and anabolic precursor molecules at the required higher rate, but using less enzyme than the catabolic pathway it replaces [8, 10–12]. Because acetate may be used by other downstream microbes as nutrient, their metabolic strategy may result from a selfish decision of the acetate-producer to invest its biosynthetic resources in this way.

The formation of byproducts not only influences the consuming microbe by offering it a potential nutrient, but it may also affect the producer. Many of these products inhibit the growth of the producer [13–16]. Acetate, ethanol and lactate are well known examples of this, as they either acidify the medium, reduce cellular (membrane) integrity, or reduce metabolic activity in another way [17]. This suggests that a second, byproduct-consuming microbe can relieve the producer of this inhibition, making cross-feeding beneficial to both species [18–20]. An extreme case of this is obligate syntrophy. Here, the producer can only grow as long as the consumer keeps the product concentration sufficiently low, as otherwise the producer’s catabolism relaxes to a state of thermodynamic equilibrium with a net zero growth rate [20]. It then only makes sense for the producer to make the byproduct when the consumer is present. This is an example in which the choice of metabolic pathway of the producer depends on the presence of a consumer. Such phenomena of consumer-affected pathway selection (obligate or not) by a producer are a main focus of this paper.

Optimal allocation of limited biosynthetic resources to maximise growth rate is an established framework to predict pathway choices for a single species [9, 21, 22]. Models in this framework may differ in the level of biological detail they incorporate; in addition to stoichiometry, reaction kinetics can also be included. Currently, microbial communities are typically modelled by combining genome-scale stoichiometric models (GEMs) [23–27], excluding kinetics. These models predict metabolic interactions and abundances of species from microbial-community extensions of flux balance analysis. They do not describe metabolite concentrations and kinetic effects. On the other end of the scale, consumer-resource models provide an alternative method that does include this [28, 29]. These models lack a biochemically consistent description of metabolism, and generally do not incorporate the concept of finite biosynthetic resources.

Here, we build a bridge between the two model extremes, by coupling biochemically consistent, coarse-grained models of metabolism and cell growth, to the dynamic and kinetic exchange of metabolites through the shared environment. We view the community as consisting of microbes that each maximise their own growth rate by optimal allocation of their biosynthetic resources over their feasible metabolic strategies in response to the current state of the environment, including nutrient and product concentrations (cf. Figure 1). In this manner, the optimal pathway chosen by a single microbe depends on the behaviour of all other species. The final community and the interactions they facilitate then form the best compromise between the metabolic potentials of all species, much like a Nash equilibrium in game theory.

**Figure 1.**
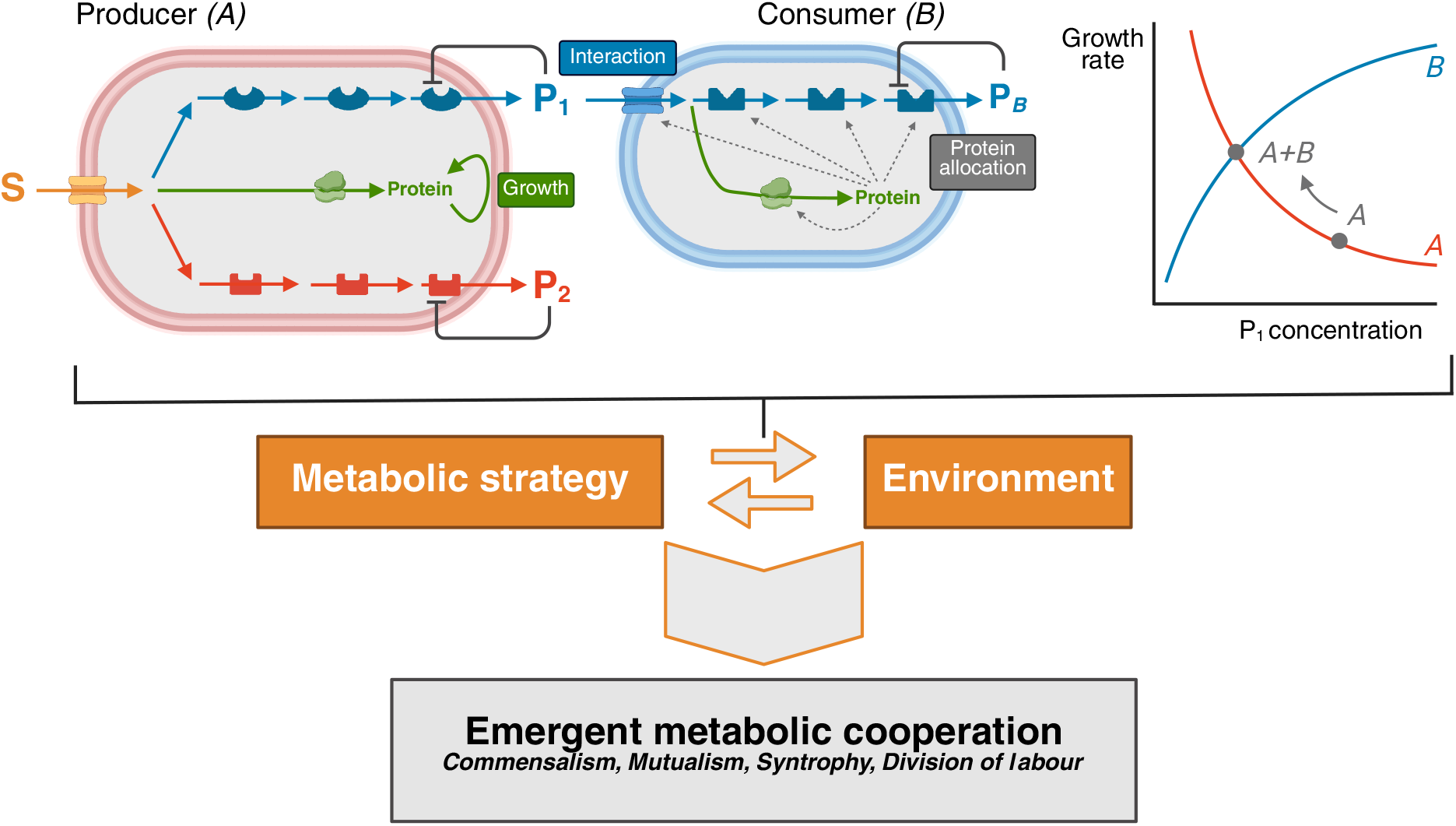
Cooperative ecological phenomena arise from interactions between microorganisms and their environment. Microbes run cellular processes by expressing proteins that catalyse the reactions in their metabolic pathways. Excretion of metabolic byproducts changes the environment, which in turn a!ects the metabolic strategy and the growth rate, for instance through product inhibition. Consumption of such a byproduct by other species then decreases its concentration, which increases the growth rate for both species. Such positive metabolic interactions can lead to emergent cooperative behaviour.

The goals of this paper are twofold: to extend the well-known resource allocation framework to multiple species, and to illustrate its approach in capturing well-known positive ecological interactions. For the latter, we will see that it is important that kinetics are incorporated. To allow the reader to get a good understanding of the structure of the optimisation problem that results, and of the influence of different modelling ingredients, we focus on communities for two and three species. In line with the single-species framework, we exclusively consider steady-state conditions. To keep the models tractable, metabolism is coarse-grained and includes kinetics. Species interact via metabolite cross-feeding, and each species is assumed to allocate a limited amount of biosynthetic resources over their feasible pathways. Thus, we view cells as selfish growthrate optimisers, and do not impose a community (fitness) objective. However, because the growth of one species impacts another by metabolite exchange, the optimisation problems are coupled. We formulate the framework such that the community composition is a result of the optimisation, instead of dictating it in advance.

The metabolic flexibility of microbes is due in large part to the volatility of their environment. However, because they are flexible, they are in fact partly responsible for this volatility, posing challenges for other species around them as well. Here we show how constrained resource allocation combined with the evolutionary compass of high growth rate allows us to make predictions on the choice of metabolic strategy of individual species and on their relative abundance in the community. We show that many positive ecological interactions may be explained by kinetic (rather than purely stoichiometric) effects in one overarching framework. This paper thus opens the door to studying larger, more complex communities whose composition is chiefly determined by metabolite interdependencies.

## 2 Results

### 2.1 Introducing the modelling approach with commensalism

In this paper, we consistently use the following terminology: a producer (*A*) grows on an initial substrate S and, as a result of its metabolism, excretes a byproduct. This byproduct is used by a consumer (*B*) to grow, thereby producing other products. The intermediary metabolite shared by the producer and consumer is consistently called the byproduct and is denoted by P_1_ or P_2_. In the models and figures, we denote producer and consumer using *A* and *B* for brevity. A second consumer is sometimes considered, and denoted with *C*.

To model metabolic interactions in a generic manner, we consider a microbial community consisting of a producer and a consumer interacting through a byproduct metabolite P_1_. We represent both species by a coarse-grained metabolic network that includes different metabolic processes, as depicted in Figure 2. The producer imports a carbon and energy source S, and catabolises it into the byproduct to generate ATP. The ATP then powers the production of biomass from the remaining carbon. The consumer grows on the byproduct using a similar metabolic network. Section 2.5 specifies the full set of assumptions and mathematical detail.

**Figure 2.**
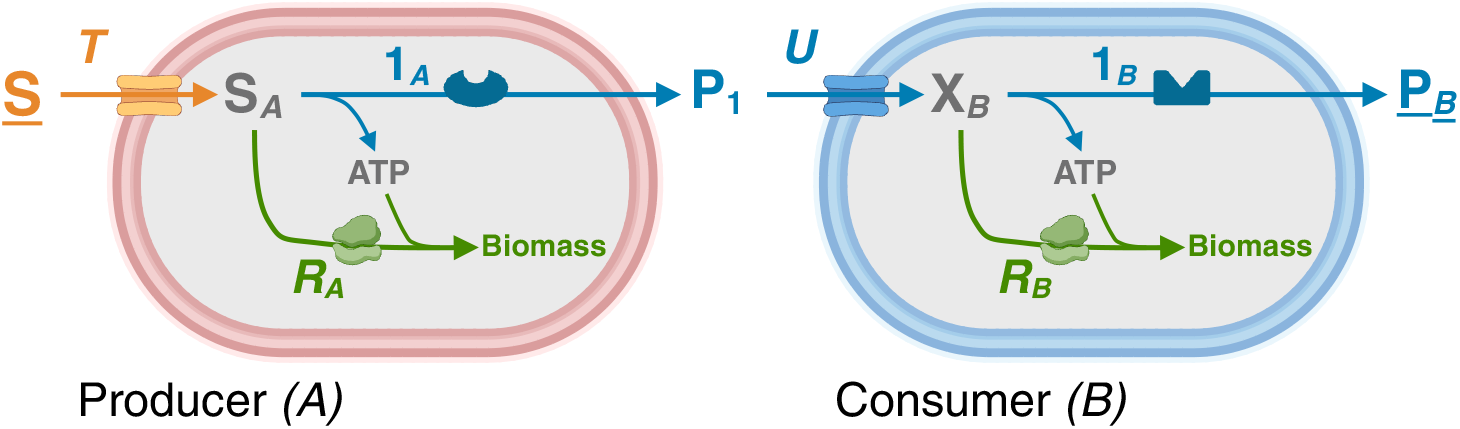
Metabolic cross-feeding is an example of commensalism. Two microbial species, a producer *A* and a consumer *B*, are represented by a coarse-grained metabolic network. The producer excretes a catabolic byproduct P_1_ that is consumed by the consumer. The consumer does not a!ect the producer’s growth in any way, but relies on the producer’s presence to provide nutrients for growth.

We include enzyme kinetics to make it possible for metabolites in the environment to influence microbial growth, not only as nutrients but also as inhibitors. In particular, as we shall see in the next section, it is important that producers are sensitive to the byproduct they excrete.

Throughout the paper we focus on balanced growth and consider steady-state metabolic activities. Each species aims to maximise its specific growth rate – the biomass synthesis rate per unit of biomass, see Section 2.5 – by optimally allocating a limited amount of biosynthetic resources over protein synthesis. All species in a stable community must have the same growth rate, which may then be referred to as the *community growth rate*: if growth rates of species would differ, one species would outgrow the other [24], violating the balanced growth condition. The species are coupled in two ways: (i) through the exchange of the metabolite P_1_ and (ii) by demanding both species grow at the same rate. A change in one of the species’ kinetic or metabolic properties is therefore expected to affect the microbial interaction, and thus also the community growth rate.

Let us now focus on the simplest example of a positive ecological interaction between two species: commensalism. In this interaction, only the consumer benefits, without impacting the producer. An example is a *Nitrobacter* consuming the nitrite produced by *Nitrosomonas* during nitrification [30].

In commensal interactions, the growth rate of the producer remains constant for any change in the properties or abundance of the consumer. In fact, even the absence of the consumer has no effect on the producer. (The “community” then consists of just the producer of course, so the community growth rate is trivially equal to the producer’s growth rate.) Obviously, the consumer can only thrive if the producer is present, because it needs the byproduct to grow. Within the model, commensalism occurs only (and always) when the producer is not sensitive to the byproduct it excretes.

Commensal interactions result in a clear unidirectional hierarchy in which species higher up in the food chain dictate the behaviour of those at lower levels. In the rest of this work, we will explore other phenomena in which the interaction is bidirectional and the consumer does affect the producer, breaking this direct hierarchy.

### 2.2 Mutualism: product inhibition induces mutually beneficial behaviour

In the case that the consumer has a positive effect on the producer, their interaction becomes reciprocal and cooperative. Such cooperative ecological interactions are referred to as mutualism. This type of interaction may occur when the byproduct inhibits the growth of the producer that excreted it. This is made possible by our inclusion of reaction kinetics in the model, i.e., the dependence of reactions on substrates and products. Product inhibition may then decrease the excretion rate of the byproduct, which, through metabolic coupling within the cell, reduces its growth rate. Such mutualistic relationships appear, for instance, in the human gut, where lactate produced by *Lactobacillus acidophilus* is consumed by *Veillonella dispar*, that enhances growth [31].

To analyse the beneficial effect quantitatively, we consider two different microbial populations (Figures 3A and B). They grow at a fixed substrate concentration (denoted by the underline, i.e. *<u>S</u>*). The first population consists of only the producer. Here, the product P_1_ accumulates in the surroundings until it reaches a steady-state concentration when excretion is balanced by diffusion into the medium. (We consider diffusion to be an implicit effect that is not modelled explicitly.) Now consider a community that contains a consumer and a producer. For this system, the steady-state concentration of P_1_ is also determined by excretion and diffusion, but additionally by uptake by the consumer. The uptake by the consumer always results in a lower concentration of P_1_. The community with the consumer thus attains a higher community growth rate (provided that the consumer can indeed attain this growth rate) compared to the community without the consumer, as indicated in Figure 3C. In other words, a producer and consumer in concert will generally grow faster than the producer alone. The exact model implementation to demonstrate this effect is discussed in Section 2.5 and Supplementary Section S10.

**Figure 3.**
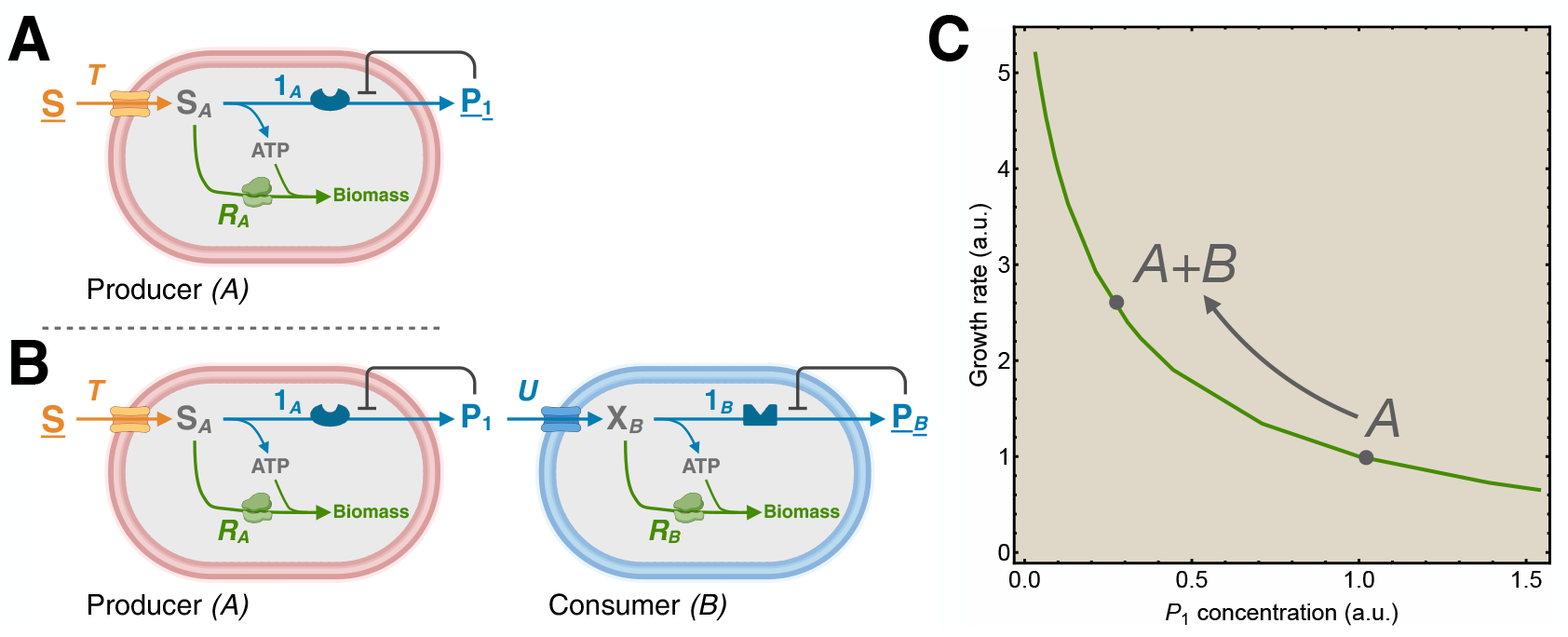
The community growth rate increases due to mutualistic cross-feeding interactions. We consider two microbial populations, **A**. one consisting of a producer in isolation and **B**. one consisting of a consumer and a producer. In both populations, product concentrations inhibit catabolic fluxes. For clarity, we only depict this for the end products P of catabolic pathways. **C**. The consumer decreases the concentration of P_1_, resulting in a reduced inhibitory e!ect. This yields a higher growth rate for the coexisting producer and consumer than for the producer in isolation. The growth rate and the concentration of P_1_ are normalised relative to the population containing only the producer.

### 2.3 Syntrophy: mutualistic interactions where coexisting species show different metabolic behaviour than species in isolation

Figure 3C indicates that the producer can also operate its catabolic pathway and grow (although at a lower rate) when the consumer is absent. However, in some cases, both species depend on each other to perform a particular process, or even to grow. This special type of mutualistic interaction is called syntrophy. We will discuss two different kinds of syntrophy: obligate and facultative.

For the first, we consider the same two populations from Figures 3A and B, consisting of only the producer or of producer and consumer together. We also did this in the previous section, but now we consider the case where the producer grows close to thermodynamic equilibrium. An example is anaerobic digestion where the hydrogen produced by acetate oxidisers is consumed by methanogens [20, 32, 33]. If the concentration of the byproduct, in absence of the consumer, now rises sufficiently, the catabolic pathway of the producer may relax to an equilibrium state of zero flux, and the net production rate of the byproduct, together with the producer’s growth rate, drops to zero, see Figure 4.

**Figure 4.**
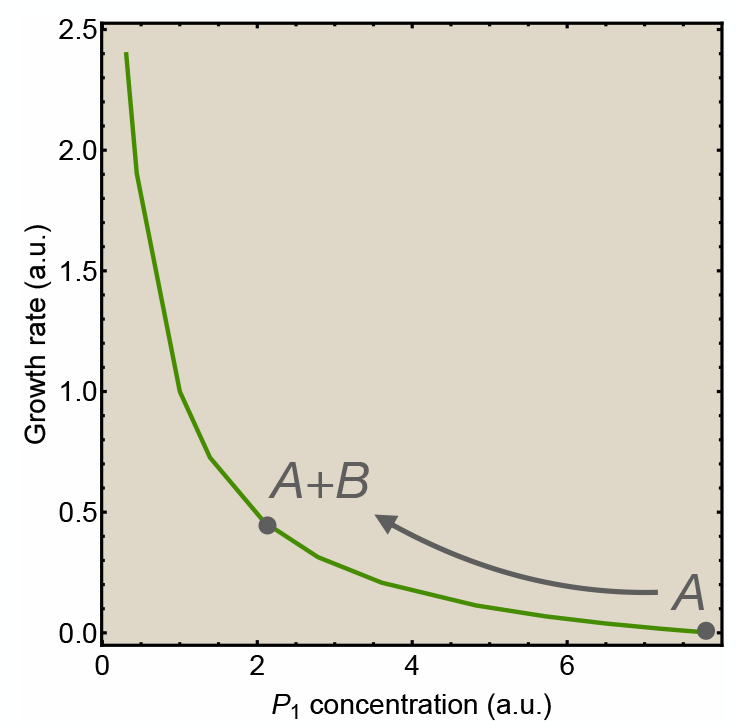
Mutualistic interactions close to thermodynamic equilibrium give rise to obligate syntrophy. Close to thermodynamic equilibrium, both species need each other’s presence to attain a nonzero growth rate. This makes their mutualistic interaction obligate. The growth rate and the concentration of P_1_ are normalised relative to the population containing only the producer from Figure 3C.

When the consumer is present, however, the additional consumption of the byproduct results in a steady-state concentration that is below its equilibrium concentration and the producer displays a net production flux of the product. Consequently, this community can attain a steady state and a nonzero growth rate. This type of syntrophy is obligate: growth of one species using a specific catabolic strategy requires the presence of the other species. In this environment, the producer can only exploit a single catabolic pathway.

The second type of syntrophy is facultative. In this case, the producer does not necessarily depend on the consumer for growth, but has alternative strategies. We represent this in our model by introducing another catabolic pathway (2_*A*_) in the metabolic network of the producer. This catabolic pathway has a different byproduct P_2_ that is not consumed by another species. The producer now uses one or a combination of its catabolic pathways to generate ATP for biomass synthesis. Existing theory [34] for single species predicts that the metabolic subnetwork that maximises the growth rate of the producer uses only one of these catabolic pathways when growth is limited by only one protein-expression constraint. For the moment, we restrict our analysis to this regime. Later, in Sections 2.4 and 2.6, we extend this to the regime where the protein expression constraints lead to the use of a combination of both catabolic pathways. A detailed analysis of the constraints and solutions of the model is given in Supplementary Sections S4 and S5.

We now show that the catabolic pathway that the producer uses to maximise its growth rate may change due to the presence of a consumer. First, we consider only the producer in Figure 5A. We assume that it maximises its growth rate using catabolic pathway 2_*A*_. We contrast this with a community in Figure 5B, consisting of a producer using catabolic pathway 1_*A*_ and the consumer of its byproduct. Although the producer on its own can sustain a lower growth rate using pathway 1_*A*_, in the presence of the consumer, however, the phenotype of the producer that uses pathway 1_*A*_ achieves a higher growth rate than the producer alone using pathway 2_*A*_. The consumer is able to decrease the steady-state concentration of P_1_ sufficiently to make the suboptimal phenotype of the producer an optimal one.

**Figure 5.**
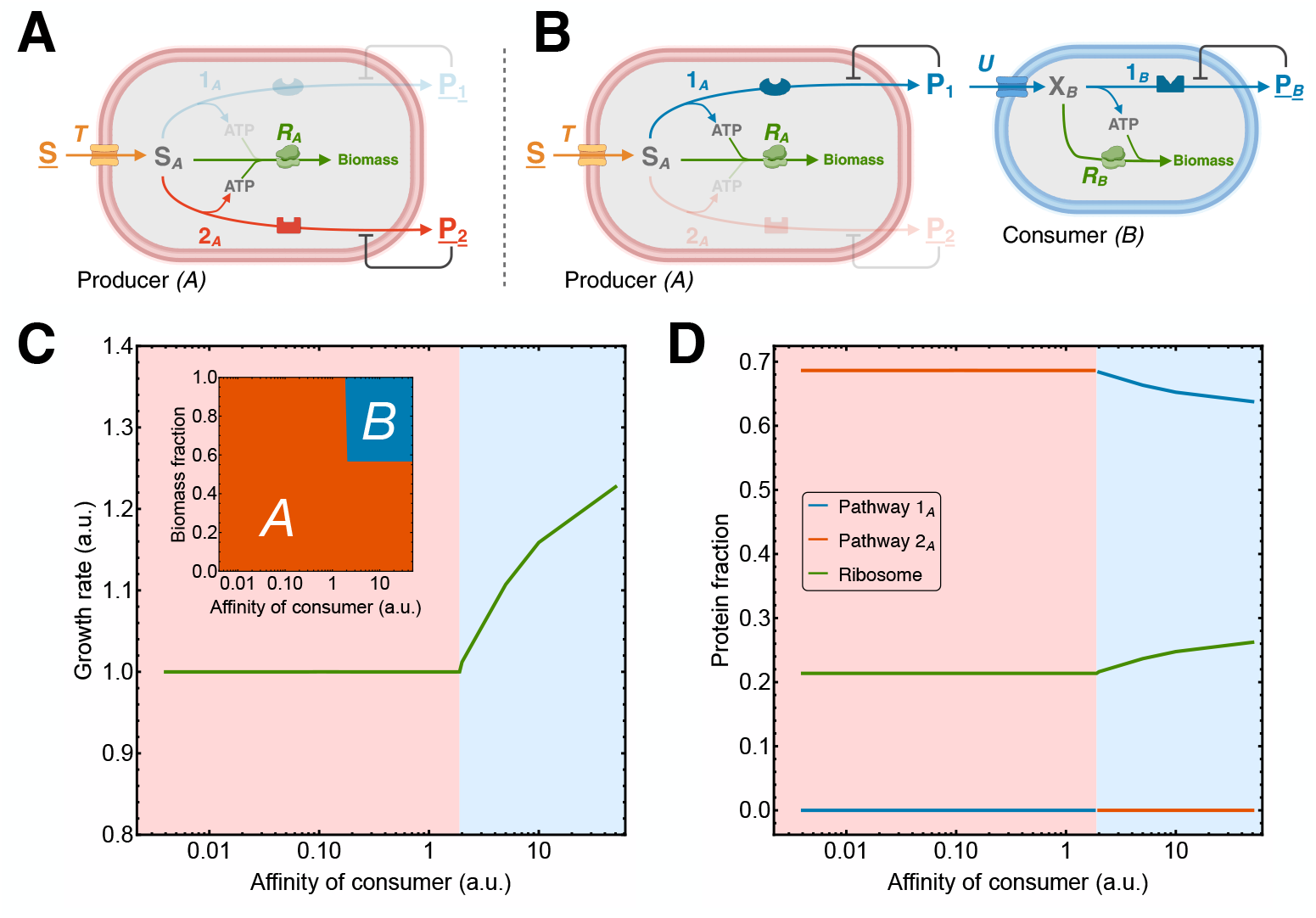
Mutualistic cross-feeding interactions lead to a switch in the optimal catabolic pathway when the producer has alternative metabolic strategies. The producer can use two di!erent catabolic pathways, 1_*A*_ and 2_*A*_, each with their own byproduct that inhibits the catabolic flux. **A**. The population containing only the producer uses catabolic pathway 2_*A*_. **B**. The population containing both the producer and the consumer uses mode 1. **C**. A discrete switch from catabolic pathway 2_*A*_ (red region) to mode 1 (blue region) occurs as the affinity of the consumer for its carbon substrate P_1_ increases beyond a threshold value. This results in an increase in the growth rate (normalised relative to the population containing only the producer), a switch in the biomass composition of the community and **D**. a switch in the protein allocation in the producer.

This type of cooperation may appear more easily when the consumer is better adapted to growing on the byproduct. When the consumer has a higher affinity for this carbon substrate, the steady-state concentration of the byproduct drops further, resulting in a higher growth rate for both species. Facultative syntrophy may thus arise when the consumer’s affinity crosses a threshold value, see Figure 5C. At this critical ‘cooperation point’, a stable community emerges in which the presence of the consumer causes a switch in the optimal catabolic strategy of the producer. The small panel in Figure 5C depicts the biomass composition of the optimal community, that also changes at the threshold affinity. Figure 5D demonstrates this switch in terms of the relative protein investment of the producer in its catabolic strategies. See Supplementary Section S11 for full detail.

An example of such facultative syntrophy occurs in mixed cultures of *Ruminococcus albus* and hydrogen-consuming bacteria that appear in ruminants. In isolation, *R. albus* ferments glucose to acetate, ethanol and some hydrogen, while in the presence of hydrogen-consuming bacteria no ethanol is produced [35, 36]. Without hydrogen consumption by these bacteria, ethanol production is required to attain redox balance.

In both cases of syntrophy discussed here, mutualistic interactions give rise to a community containing two species that exhibit different metabolic behaviours than they would in isolation.

### 2.4 Division of labour as a special case of facultative syntrophy

In the previous section we considered a metabolic switch between two catabolic strategies due to a syntrophic interaction. We can examine division of labour (DOL) in a similar fashion. Giri et al. [37] established general criteria for classifying microbial interactions as DOL and provided examples of this phenomenon. Here, we restrict our analysis to scenarios in which the degradation of a substrate to a final product is carried out by either one species, or by two coexisting species through cross-feeding. The latter case is termed a DOL strategy. Contrary to mutualism and syntrophy, in DOL it is the same overall conversion of substrate to final product that is carried out by one or by two species. To give one example, the bacterium *Pseudomonas stutzeri* either completely degrades nitrate to nitrogen gas by itself, or cross-feeds the intermediate nitrite to a second microbe [38], which in turn produces the nitrogen gas.

DOL can be incorporated into our model formalism by assigning specific properties to the catabolic pathways of the producer and the consumer, which are depicted in Figures 6A and B. Mode 1 now represents the DOL pathway with cross-feeding, in which the producer catabolises the substrate S into the byproduct P_1_, that is further degraded by the consumer to the end product P_2_. Mode 2 represents the complete degradation of the substrate to the end product P_2_, performed by the producer alone. In this scenario, there is a fundamental tradeoff that microbes need to overcome. A longer pathway has a higher Gibbs free energy potential, which may be used for the production of more ATP. Longer pathways thus allow increasing the ATP yield compared to the shorter pathway that only partially degrades the substrate. However, a longer pathway requires investing limited enzyme resources into more individual reactions, thus lowering the amount they can invest in each step. This reduces the pathway flux. Whether higher growth rates are achieved through a high ATP yield or a high catabolic rate decides whether a single species or a two-species system is favoured. The details of the model are fully described in Supplementary Section S12.

**Figure 6.**
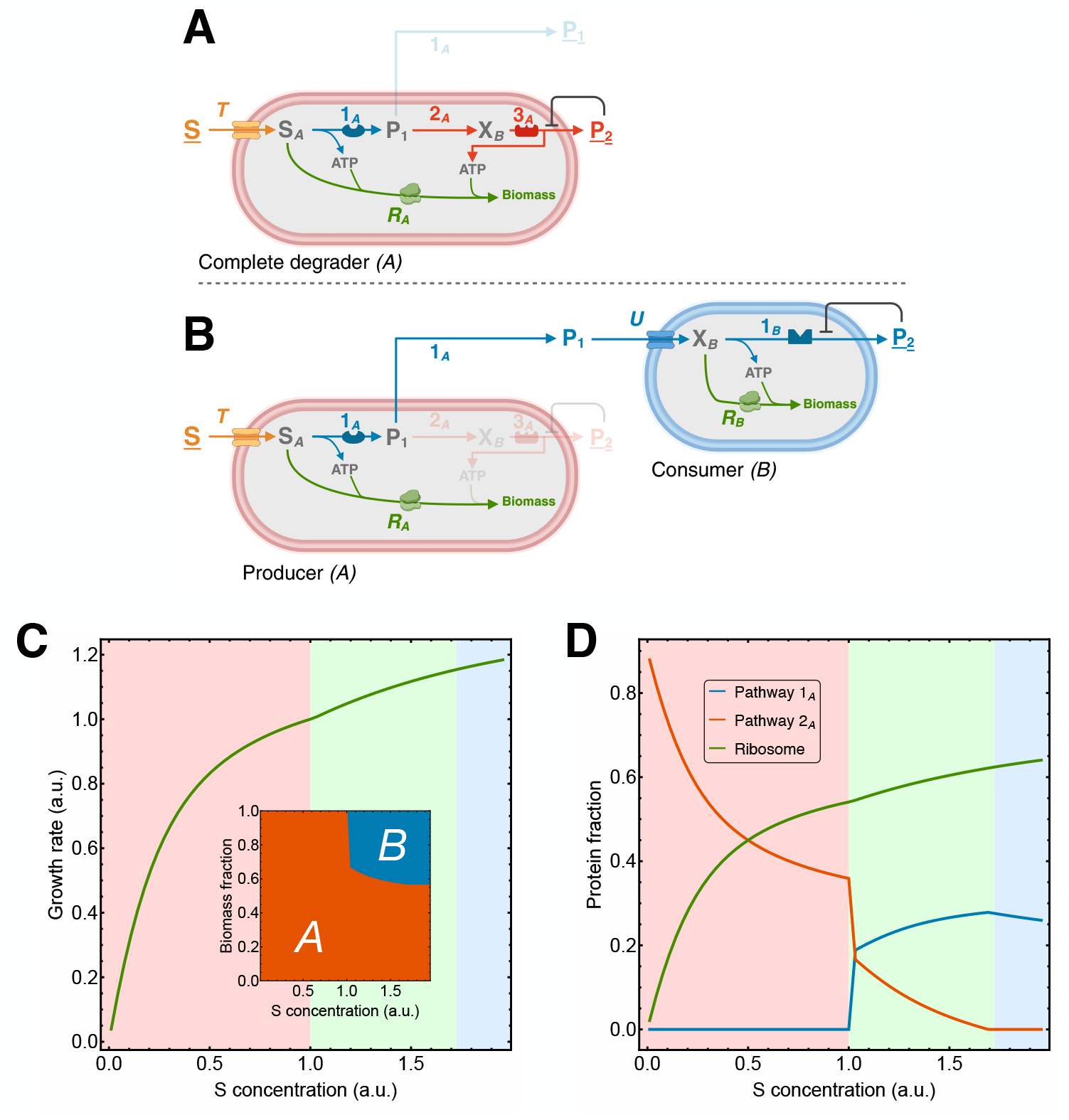
A shift from complete degradation to division of labour occurs for increasing substrate concentration. **A**. The population containing only the producer performs the complete degradation of the substrate S to the end product P_2_. **B**. In the two-species community, the last two reactions of this degradation pathway are taken over by a consumer of the intermediate P_1_. The reactions in the consumer describe the same chemical conversions, but with di!erent reaction-kinetic properties. **C**. A gradual shift from complete degradation by the producer to DOL occurs as the substrate concentration *S* increases beyond a critical value. This shift appears in the biomass composition of the community and **D**. in the protein allocation in the producer. In the red region complete degradation by the producer is optimal, in the green region a combination of both strategies is optimal and in the blue region DOL is optimal. The growth rate and the substrate concentration are normalised relative to the critical point where the shift initiates.

Similar to facultative syntrophy in Figures 5C and D, for DOL we observe a discrete switch in the optimal catabolic strategy from mode 2 to mode 1 when the affinity of the consumer for P_1_ increases beyond a critical value (see Figure S8 in Supplementary Section S12). This represents a switch from a complete pathway in a single species to a DOL strategy in a community of two species. In other words, our model predicts that DOL may arise from the presence of the consumer, provided that it has a sufficiently positive effect on the producer by decreasing the steady-state concentration of P_1_.

In most studies, DOL is not studied from a product perspective as we have done here, but rather from the effect of the substrate concentration on the producer. Complete (long) pathways with higher biomass yield are often found at low substrate concentrations, while DOL strategies are found at higher substrate concentrations, presumably leading to faster growth [39–41]. Our model is consistent with these observations, as illustrated by Figures 6C and D. These figures show a shift in protein investment from mode 2 to mode 1 in the producer as *S* increases beyond a critical value. Note that the shift is gradual instead of switch-like. This is explained in more detail in Supplementary Sections S6 and S12. The small panel in Figure 6C illustrates that this shift enables growth of the consumer, as its carbon substrate P_1_ is now produced. But the shift also happens in absence of the consumer. It is entirely caused by the constraints on protein investment in the producer. Nevertheless, the new byproduct that results from the combination of metabolic modes in the producer does allow the consumer to take advantage, and convert the new byproduct to a final end product, so we can still term this DOL.

### 2.5 Modelling microbial communities as a coupled optimisation problem

#### Construction of the coarse-grained reaction network for two species

We provide a coarse-grained description of microbial growth, either of a single producer, or of a producer-consumer pair that interacts via an intermediate byproduct metabolite. Each species is equipped with a coarse-grained metabolic network, depicted in Figures 5A and B, and consists of different metabolic processes, similar to the core model used in Droste et al. [12]. Metabolic fluxes are denoted by *v* with subscripts denoting the individual processes. The producer imports a carbon and energy source S via a transport reaction (*T*, so with flux *v*_*T*_). The producer has two ways to catabolise this substrate; it can use them combined or separately. The catabolic pathway 1_*A*_ converts the internal carbon S_*A*_ into 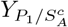 of product P_1_ and produces 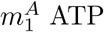 per S. The catabolic pathway 2_*A*_ converts S_*A*_ into 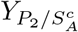 of product P_2_ and produces 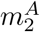 ATP per S. The remaining carbon is used to synthesise biomass by consuming 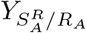 S_*A*_ and 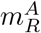 ATP. We model biomass formation solely as protein synthesis by the ribosome (*R*_*A*_), since this process consumes most ATP during growth. Consequently, the biomass concentration is represented by the total protein concentration 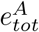 in the cell. The consumer is modelled in a similar fashion, except that uptake (*U*) of its carbon source P_1_ describes both its transport and conversion into 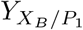 of intermediate X_*B*_. The reason for this minor modification is explained in Supplementary Section S4. The intermediate X_*B*_ is catabolised by pathway 1_*B*_, which converts it into 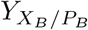 of product P_*B*_ and produces 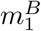 ATP per X_*B*_.

The concentrations of the external metabolites (S, P_2_, and P_*B*_) are assumed to be fixed, which is denoted by the underline in Figures 5A and B, i.e., *<u>S</u>*. The concentrations of the internal carbon intermediates and ATP in both species are described by the following set of differential equations,

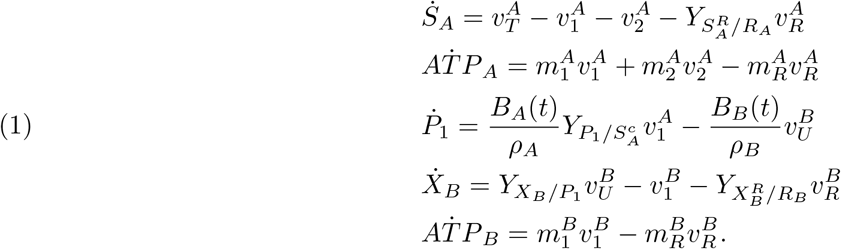

Here, *B*_*A*_(*t*) and *B*_*B*_(*t*) are the biomasses (in gram dry weight per litre of culture vessel) of both species and *ω*_*A*_ and *ω*_*B*_ are their densities (in gram dry weight per litre cell), that are assumed to be constant. We denote the stoichiometric matrix following from these equations by **N** and the corresponding flux vector by

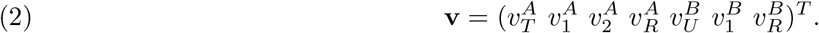

In balanced growth, all concentrations are in steady state [42]. This holds for the internal metabolic concentrations but also for exchange metabolite P_1_ in the medium. Together, these steady state equations are summarised by **Nv** = **0**.

To model the rates explicitly, we represent each metabolic process in Figures 5A and B by a biochemical reaction catalysed by its specific enzyme. Each enzymatic reaction *j* has a corresponding enzyme concentration *e*_*j*_, catalytic rate constant *k*_*j*_ and runs at rate *v*_*j*_ = *k*_*j*_*e*_*j*_*f*_*j*_(**x**). The saturation factor *f*_*j*_ describes the saturation of the enzyme; we assume it is constant for biomass synthesis in both species. For the reactions representing catabolic processes, we include the concentrations (**x**) of the carbon substrates and products. The corresponding saturation factor follows reversible Michaelis-Menten kinetics [43]. This explicitly captures both the thermodynamic and kinetic product inhibition effects.

#### Optimisation of the community growth rate

During balanced growth at steady-state growth rate *ε* all intracellular concentrations remain constant. The total protein concentration *e*_*tot*_, representing the biomass in our case, is generally observed to be constant across conditions [44–46]. This implies that the protein synthesis rate by ribosomes *v*_*R*_ is balanced by the rate of dilution by growth 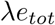 [47],

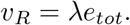

This yields an expression for the growth rate

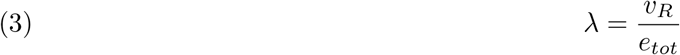

as the protein synthesis rate per unit of protein. We assume that both species maximise their steady-state growth rate, respectively 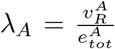 and 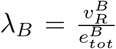 as this gives the highest fitness in constant conditions [48].

In the previous sections, we have considered populations that consist of either a producer in isolation or a producer-consumer pair. We want to formulate the growth rate maximisation problem such that it describes both these situations, meaning that the growth-rate-maximising community may consist of two species, or of only one species, in which case the “community” is a monoculture. We therefore model the coupling of the species as facultative rather than hard-coded. This metabolic coupling is described by the differential equation for the concentration of the exchange metabolite P_1_ in eq. (1). In the case that both species are present and they exchange P_1_, their biomass increases due to growth. In Supplementary Section S3 we show that they necessarily grow at an equal rate *λ*_*A*_ = *λε*_*B*_ to ensure steady-state for P_1_. Here, we also derive that the biomasses of both species are coupled via

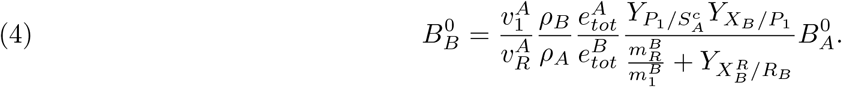

This relationship determines the steady-state biomass of the consumer 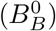 in terms of the biomass of the producer 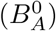. It shows that excretion of P_1_ at rate 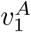 always leads to the presence of the consumer in this model, except when the P_1_ concentration is fixed, such as in Figure 3A. When the producer allocates less protein to mode 1, the flux ratio 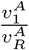 decreases and less of the byproduct P_1_ is excreted. Since all other parameters on the right-hand side are constant, this implies a decrease in the steady-state biomass of the consumer.

We note that this relationship remains true even when the consumer is not present. When the producer uses pathway 2_*A*_ exclusively, the steady-state excretion flux 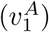 of P_1_ is zero. This means that there is no exchange of P_1_ and it removes the carbon source for the consumer, so it will not synthesise any biomass. This situation is captured by eq. (4) when both sides are equal to zero. This relationship thus describes the facultative coupling of the species. Including this relation is essential to model a steady-state of the two-species community, but it naturally vanishes in the case where no consumer is present. This relationship is therefore a more general condition for the community steady-state than equality of the growth rates, as it holds irrespective of the number of species in the community.

We now formulate maximisation of the (community) growth rate as an optimisation problem by

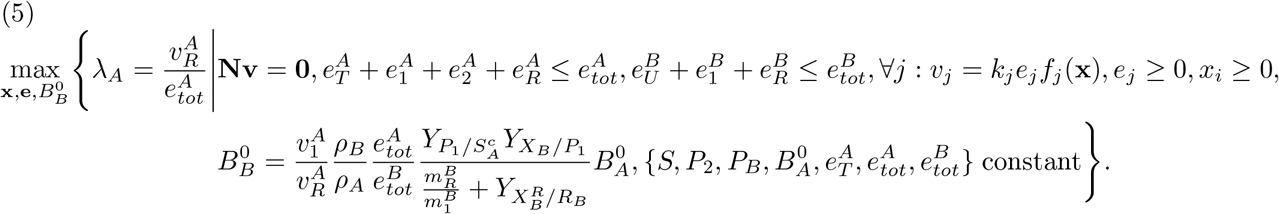

We pick the growth rate of the producer as the objective function, which describes the (community) growth rate in both the one- and two-species populations. The optimisation variables are the internal metabolite concentrations **x** = *{S*_*A*_, *P*_1_, *X*_*B*_*}*, the enzyme concentrations 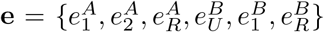 and the biomass 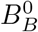 of the consumer. Because the relation in eq. (4) holds with and without a consumer present, this optimisation problem describes both populations consisting of either one or two species. Accordingly, the model does not dictate the community composition, but rather infers it as an outcome of the optimisation.

The optimisation problem and its solutions are studied in more detail in Supplementary Section S5. In Supplementary Sections S9-S12, we explain how the phenomena discussed in Sections 2.2-2.4 are represented by the general model in terms of a single species vs a two-species microbial community. For all these variants of the general model, a Mathematica implementation is provided in the source code on Zenodo.

### 2.6 Example applications

In this section we use our modelling framework to derive predictions for positive interactions that are different from the previously discussed phenomena, but that can also be tested experimentally.

#### Syntrophic interactions with multiple consumers may determine the optimal exchange metabolite

For the first example, we consider the situation for facultative syntrophy as presented in Figures 5A and B. We extend the single species population with another species *C* that consumes the product (P_2_) of catabolic pathway 2_*A*_, resulting in the community shown in Figure 7A. The additional consumption of P_2_ by species *C* leads to a steady-state concentration that is lower than that achieved (implicitly) by diffusion alone. The growth rate of the community with species *A* and *C* thus exceeds that of the population with only species *A* from Figure 5A, as the red region in Figure 7C illustrates, due to reduced product inhibition.

**Figure 7.**
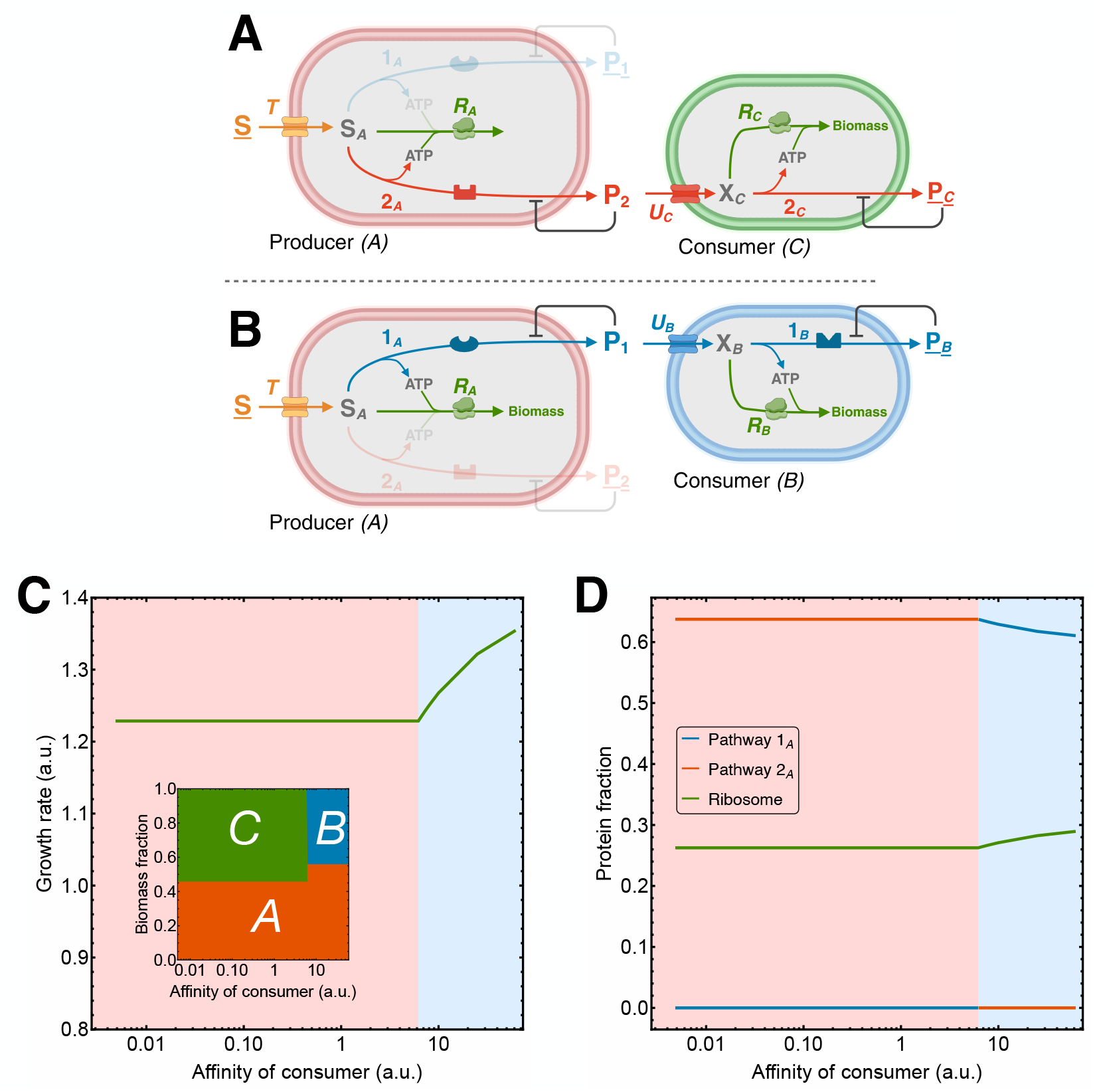
Switch in the optimal catabolic pathway for syntrophic interactions with multiple consumers. **A**. A community consisting of the producer and consuming species *C* uses catabolic mode 2 to exchange P_2_ and degrade it further into P_*C*_. **B**. The community containing the producer and consumer *B* uses catabolic mode 1 to exchange P_1_ and degrade it into P_*B*_. **C**. A switch from catabolic pathway 2 (red region) to mode 1 (blue region) occurs as the affinity of the consumer for P_1_ increases beyond a critical value. This results in an increase in the growth rate (normalised relative to the growth rate of the population containing only the producer from Figure 5C), a switch in the biomass composition of the community, and **D**. a switch in the protein allocation in the producer. The critical affinity at which the switch occurs is higher than in Figure 5C due to consumption of P_2_ by species *C*.

For the community in Figure 7B, consisting of the producer and consumer *B*, to achieve an even higher growth rate, this means that *B* should lower the steady-state concentration of P_1_ even more. This implies that the critical affinity of *B* for P_1_, at which both communities grow equally fast, increases compared to its value in Figure 5C. Again, beyond this critical cooperation point, a switch in the optimal catabolic pathway of the producer occurs, as indicated in Figure 7D. The condition for this switch has become more restrictive, but it remains qualitatively similar to the results for facultative syntrophy in Figures 5C and D. If *P*_1_ reaches zero before or at the critical cooperation point, the community with *B* as consumer cannot achieve a higher growth rate than the one in which *C* is the consumer.

The characteristics of consumers *B* and *C* thus determine the optimal behaviour of the producer, in particular the byproduct that the producer should excrete to achieve the highest growth rate. Suppose that consumers *B* and *C* are different phenotypes of the same species, producing the same byproduct (P_*B*_ = P_*C*_) but growing on a different carbon source supplied by the producer. Using our model, we can compute which of these consumer-phenotypes and corresponding cross-fed metabolite will result in the highest community growth rate. This may explain why the cross-feeding of one specific metabolite is more beneficial for the community than another metabolite. In other words, in this way we can predict the optimal exchange metabolite for the producer, and how this depends on the consuming species.

#### Resource allocation over mixed strategies changes due to catabolic product consumption

For the next illustration, we turn to the regime where the producer uses a combination of both catabolic pathways to maximise its growth rate. How this regime is characterised is explained in more detail in Supplementary Sections S5 and S6.

As shown in Figure 8A, the producer on its own excretes two byproducts, whose steady-state concentrations *P*_1_ and *P*_2_ are implicitly determined by the balance between excretion and diffusion. We compare its performance to a microbial community, in which an additional consumer harvests the product P_1_ (cf. Figure 8B), thus decreasing its steady-state concentration (with respect to the population with only the producer) when its affinity increases. This reduces product inhibition for catabolic pathway 1_*A*_, which now requires less protein to attain the same flux. Therefore, it is advantageous for the producer to increase the relative protein investment in catabolic pathway 1_*A*_ compared to pathway 2_*A*_, so the remaining protein can be invested in biomass synthesis to increase the growth rate. This increased protein efficiency of mode 1 due to an increased consumer affinity thus increases both the community growth rate and the protein fraction allocated to mode 1, as shown in Figures 8C and D. Panel D shows a gradual shift in the optimal combination of strategies from mode 2 to mode 1 that is similar to the shift from complete degradation to a DOL strategy when the substrate concentration increases, as observed in Figure 6D. The regime where the producer uses a combination of catabolic pathways instead of one thus results in a change in the optimal strategy that is qualitatively different from the discrete switch observed in the other cases of syntrophy presented in Figures 5D and 7D. However, if the consumer increases its affinity for P_1_ significantly, this may induce a complete shift from mode 2 to mode 1, as Figure 8D illustrates. The corresponding model is analysed in more detail in Supplementary Section S14.

**Figure 8.**
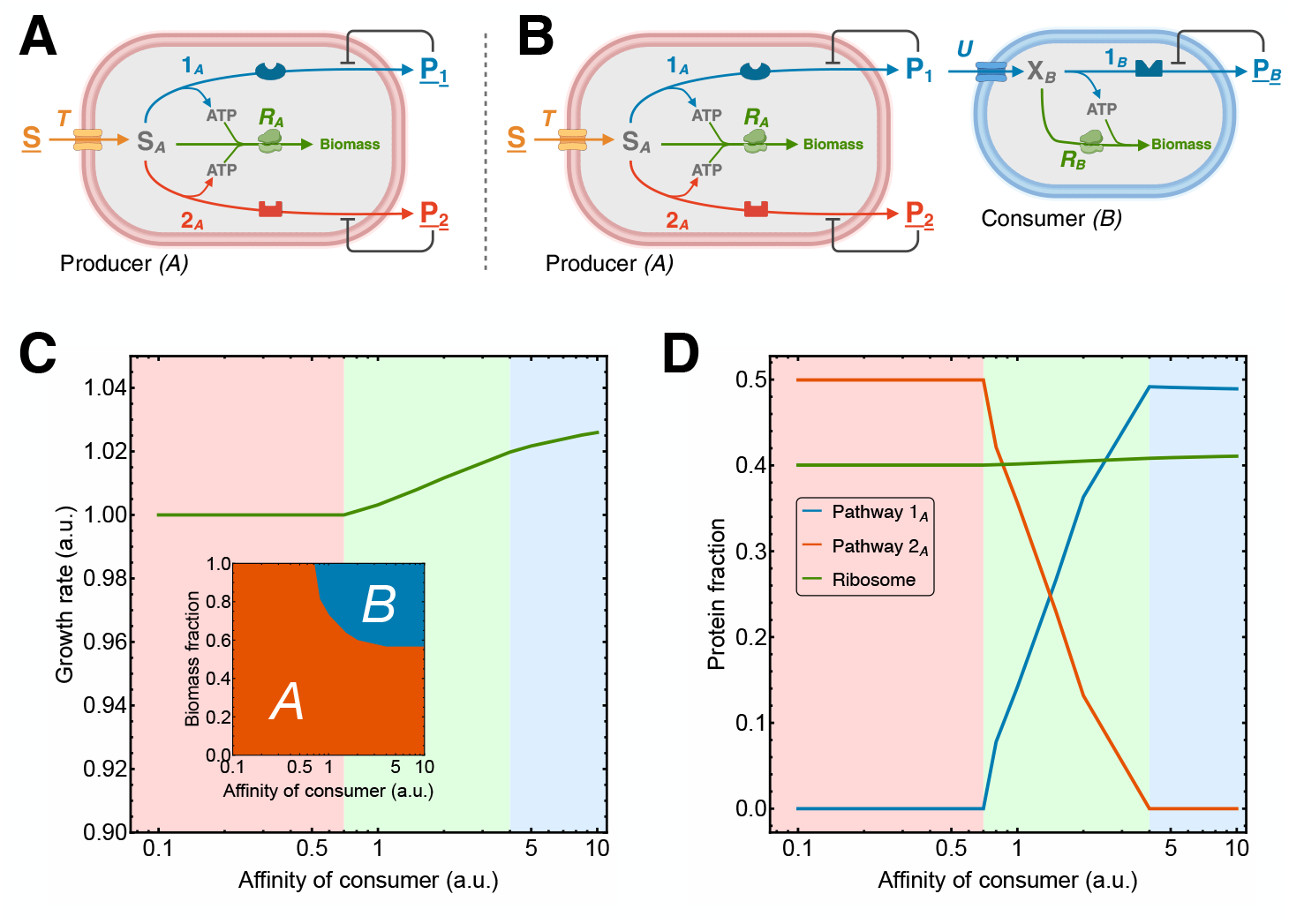
Resource allocation over both catabolic pathways changes due to byproduct consumption. **A**. In both populations the producer uses both catabolic pathways and excretes both byproducts. **B**. In the population containing both the producer and the consumer, the consumer feeds on the byproduct P_1_ of mode 1. **C**. A gradual shift from catabolic mode 2 to mode 1 occurs as the affinity of the consumer increases beyond a critical value. This shift appears in the biomass composition of the community and **D**. in the protein allocation in the producer. In the red region mode 2 is optimal, in the green region a combination of both modes is optimal and in the blue region mode 1 is optimal. The growth rate is normalised relative to the population containing only the producer.

We may thus predict when the consumer alters the amount of resources invested by the producer in its metabolic modes through consumption of one of the byproducts of the producer. This can be verified experimentally by changing the concentration of one of the byproducts of a microbial species in a bioreactor (e.g. by adding this product to the feed). Proteomic profiling of each catabolic pathway before and after this perturbation should identify alterations in the relative protein allocation among these modes (e.g., using SIP [49]).

## 3 Discussion & Conclusion

In this work, we have investigated microbial communities that involve metabolic interactions between species. Due to these interactions, the optimal metabolic pathway used by a single microbe to maximise its growth rate depends on all other species. This fact motivated our study of microbial communities. We extended the framework of optimal allocation of limited biosynthetic resources from single species to communities of species. We were able to capture mutualistic, syntrophic, and division of labour interactions using this framework. These phenomena involve cooperative interactions, which we model via the exchange of a metabolic byproduct that inhibits growth of the producer and enhances the growth of the consumer. For different types of syntrophic interactions, we demonstrated that a change in community composition coincides with either a discrete switch or gradual shift in the optimal metabolic strategies of the species.

The general framework offers a bottom-up, mechanistic, approach to study microbial communities from the perspective of individual species. We formulated this framework for two microbial species that exchange a metabolite. By coupling them in this manner, and demanding maximisation of the growth rate of the producer, we were able to find communities in which producer and consumer together reached a higher (and equal) growth rate than the producer alone. Their coupling also resulted in relation (4) that determined the relative abundances of both species, and that induced a number of additional phenomena. For instance, a higher biomass yield of the consuming species on its carbon substrate induces a higher relative biomass fraction, but it also results in a lower byproduct concentration and thus a higher growth rate for both species. More details may be found in Supplementary Section S3-S7. Such counterintuitive effects complicate reasoning about microbial communities and motivate usage of mathematical modelling methods [7, 50, 51].

Since all ecological phenomena discussed here are modelled by a metabolite cross-feeding interaction, we also created a common vocabulary that allows for comparison between these different ecological interaction types in mechanistic terms. For mutualism and obligate syntrophy, lowering the byproduct concentration by the consuming species is sufficient to give rise to a two-species community. Experimental work [52, 53] shows how evolution may give rise to these phenomena. For facultative syntrophy, this decrease should be sufficiently significant so that a critical “cooperation point” is passed and a metabolic switch occurs. This critical cooperation point depends on environmental conditions and species’ traits. We also showed that DOL may be interpreted as a special case of facultative syntrophy. Both these phenomena require that the producing species can attain comparable growth rates with both its metabolic strategies. In Supplementary Section S9 we show how each phenomenon is described by a variant of our model in a different parameter regime.

Microbial communities are often either considered to live in fixed environmental conditions, and then studied using detailed metabolic models [23–25, 27], or the interaction with their environment is put center stage, using phenomenological descriptions of cell growth [29, 54]. Here we have tried to build a bridge between these two worlds, recognising that microbes change the environment they inhabit by nutrient consumption and byproduct excretion, but using coarse-grained descriptions of metabolism to capture the importance of limited resource allocation for biosynthesis. We stress that we have explicitly included the kinetic dependence of reaction rates on metabolite concentrations. In each case, it is the relief of product inhibition for the producer by additional consumption by the consumer that gives rise to a higher growth rate for the producer (and thus also for the consumer). A purely stoichiometric model with fixed enzyme saturations cannot capture this type of synergistic effect of the consumer on the producer. Another benefit of our approach is that it does not stipulate up front whether a consumer is actually present in the optimal growth state. By setting the producer’s growth rate as objective, we have allowed for the possibility that the consumer does not grow at all, or indeed is present and then matches the producer’s growth rate. This is particularly relevant for future studies with larger communities, in which the community composition is not clear a priori and should be the result of the interactions between the community members.

Coarse-grained kinetic approaches to study cross-feeding in microbial communities similar to ours do exist. Mauri et al. [55] find conditions for coexistence in a consortium of two *E. coli* strains, while Lopez et al. [56] identify different forms of metabolite toxicity that lead to stable cross-feeding interactions. Pfeiffer and Bonhoeffer [40] and Kreft [39, 41] also used principles from optimal resource allocation to study cross-feeding and DOL, and even derive an optimal pathway length based on earlier results from Heinrich et al. [57]. Although our work focuses mainly on maximisation of the (community) growth rate, these studies are for a large part supplementary to our general framework.

Species themselves are also shaped by their environment. Evolution through environment-dependent selection may cause modifications of physiological properties such as, for instance, the transporter affinity for certain nutrients and products. We demonstrated that altering this affinity can result in changes in metabolic behaviour and community composition. This is especially interesting when substrate import occurs through facilitated diffusion. Since this is a passive process that does not require ATP, the equilibrium constant of this process equals one. The Haldane relation [58] then implies a trade-off between the affinity and the rate constant of substrate import [59]. In nutrient rich-environments, one would therefore expect species with low affinity but high rate transporters, while in poor environments, high affinity transporters with low rate constants are expected. Following the reasoning for DOL from Section 2.4, shorter pathways are preferred in rich environments, while poor environments give rise to pathways with more reactions. These theoretical findings have been observed for bacteria involved in (de)nitrification [38, 60].

We have focused on small microbial communities to aid the reader how one can build models that combine coarse-grained metabolism with dynamic nutrient environments, and apply maximisation principles that do not prescribe in advance which species will coexist. Clearly, real communities are larger and exhibit more complex interactions. This paper provides a foundation to extend the theory in that direction. For organisms that are involved in a linear degradation chain, we can use the formalism to investigate how non-nearest neighbours in the chain affect each other. We could also apply our analysis to growth of organelles in eukaryotic cells, as synthesis of a new cell requires balanced growth of all its components at the same rate. Furthermore, compartmentalisation has been shown to result in a lower product inhibition effect [61].

Overall, our modelling framework opens the door to study consortia of metabolically-coupled agents. Since our formalism is based on fundamental biochemistry, it captures different ecological phenomena in mechanistic terms.

## Supporting Information

### Supplementary Appendices

Supplementary sections S1-S14 with glossary, derivations and additional background information for the models for different cooperative ecological phenomena. (PDF)

### Source codes

Mathematica notebooks used for numerical implementations of the variants of the community model. These notebooks are used to create the graphs in all figures. Source codes are available on Zenodo (https://doi.org/10.5281/zenodo.23694644). (NB)

## Acknowledgements

We would like to thank our colleagues of the VU Systems Biology Lab for critical and stimulating discussions. MJD and RP acknowledge funding by NWO grant 613.009.131 ‘Control of maximal growth rate by single-celled organisms’. Figures containing metabolic networks are created using BioRender.

