## Supplementary Appendices for "Emergent metabolic cooperation in communities of microbes that maximise growth rate by optimal protein expression"

### S1 Glossary

*A*: Microbial species that produces and excretes a byproduct.
*B*: Microbial species that consumes the byproduct $P_1$ as carbon and energy source.
*B(t), B<sup>0</sup>*: The biomass of a microbial species, as function of time or in steady-state (gDW/L culture).
*C*: Microbial species that consumes the byproduct $P_2$ as carbon and energy source.
*e<sub>j</sub>*: Concentration of the enzyme catalysing reaction *j* (mmol protein/L cell).
*e<sub>tot</sub>*: The total concentration of protein in a cell (mmol protein/L cell).
*f<sub>j</sub>*: Saturation factor of the enzyme catalysing reaction *j*, may depend on metabolite concentrations **x** involved in the reaction.
*k<sub>j</sub>*: Catalytic rate constant of reaction *j* ($h^{-1}$).
*m<sub>R</sub>*: The amount of ATP required to synthesise one mole protein (mmol ATP/mmol protein).
*m<sub>j</sub>*: ATP yield of reaction *j* per mole of the carbon source (mol ATP/mol S).
*P*: Byproduct of a catabolic pathway that is excreted by the cell.
*S*: Carbon and energy source.
*v<sub>j</sub>*: The rate of reaction *j* (mmol/h L cell).
*X*: Carbon intermediate for consuming species.
*x<sub>i</sub>*: Concentration of metabolite *i* (mmol/L cell).
*Y<sub>P/S</sub>*: Biomass yield of product *P* per mole substrate *S* (mol P/mol S).
*Y<sub>S/R</sub>*: The amount of carbon source required to synthesise one mole of biomass (mol S/mol protein).
$\lambda$: Growth rate of a microbial culture ($h^{-1}$).
$\rho$: The biomass density of a microbial species (gDW/L cell).

### S2 Mathematical details of the coarse-grained reaction network for two species

In this section, we discuss additional features of the coarse-grained reaction network of the microbial community presented in Figure S1, for which the main modelling ingredients were explained in Section 2.5. This community consists of a producer (*A*) and a consumer (*B*) that exchange metabolite $P_1$, which is a byproduct excreted by the producer and a carbon and energy source for the consumer.

We study this community in a fixed external environment. This means that the external metabolites *S*, $P_2$ and $P_B$ have reached a steady-state concentration that is determined by molecular diffusion and through production and consumption by other surrounding species. Since these processes are not explicitly modelled for external metabolites, we fix the concentrations of these metabolites.

The concentrations of the internal metabolites (including $P_1$) are described by the set of differential equations in eq. (1). The corresponding stoichiometric matrix is given by

$$(S1) \quad \mathbf{N} = \begin{pmatrix} 1 & -1 & -1 & -Y_{S_A^R/R_A} & 0 & 0 & 0 \\ 0 & m_1^A & m_2^A & -m_R^A & 0 & 0 & 0 \\ 0 & \frac{B_A(t)}{\rho_A} Y_{P_1/S_A^c} & 0 & 0 & -\frac{B_B(t)}{\rho_B} & 0 & 0 \\ 0 & 0 & 0 & 0 & Y_{X_B/P_1} & -1 & -Y_{X_B^R/R_B} \\ 0 & 0 & 0 & 0 & 0 & m_1^B & -m_R^B \end{pmatrix}.$$

During balanced growth, the concentrations of the internal metabolites are in steady-state, which is described by $\mathbf{N}\mathbf{v} = \mathbf{0}$ where **v** is the flux vector in eq. (2).

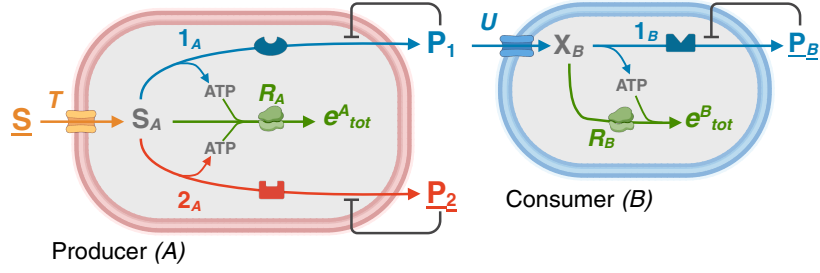

FIGURE S1. **The coarse-grained reaction network for a microbial community consisting of a producer and a consumer.** A carbon source $S$ is transported ($T$) into the producer ($A$). This microbe has two catabolic pathways, $1_A$ and $2_A$, that generate energy carriers (ATP) while converting $S$ into respective catabolic byproducts $P_1$ and $P_2$. Protein synthesis by the ribosome ($R_A$) consumes the remaining carbon and generated ATP. The consumer ($B$) is modelled by a similar metabolic network and has one catabolic pathway $1_B$ with byproduct $P_B$.

The steady-state flux vectors that correspond to the metabolic modes using exclusively catabolic pathways $1_A$ and $2_A$ are, respectively,

$$(S2) \quad \mathbf{v}_1 = v_R^A \begin{pmatrix} \frac{m_R^A}{m_1^A} + Y_{S_A/R_A} \\ \frac{m_R^A}{m_1^A} \\ 0 \\ 1 \\ \frac{m_R^A}{m_1^A} \frac{B_A(t)\rho_B}{B_B(t)\rho_A} Y_{P_1/S_A^c} \\ \frac{m_R^A}{m_1^A} \frac{B_A(t)\rho_B}{B_B(t)\rho_A} \frac{m_B^B}{m_1^B} \frac{Y_{P_1/S_A^c} Y_{X_B/P_1}}{\frac{m_B^B}{m_1^B} + Y_{X_B/R_B}} \\ \frac{m_R^A}{m_1^A} \frac{B_A(t)\rho_B}{B_B(t)\rho_A} \frac{Y_{P_1/S_A^c} Y_{X_B/P_1}}{\frac{m_B^B}{m_1^B} + Y_{X_B/R_B}} \end{pmatrix}, \quad \mathbf{v}_2 = v_R^A \begin{pmatrix} \frac{m_R^A}{m_2^A} + Y_{S_A/R_A} \\ 0 \\ \frac{m_R^A}{m_2^A} \\ 1 \\ 0 \\ 0 \\ 0 \end{pmatrix}.$$

Both these flux vectors must have positive entries and are normalised such that the biomass production flux of the producer equals 1. The second flux vector $\mathbf{v}_2$ shows that the consumer does not grow (all its fluxes are zero) when the producer uses catabolic pathway $2_A$ exclusively. This mode is only included in the models for phenomena that exhibit a switch or a shift in metabolic strategy.

The flux vectors (S2) require each reaction in the corresponding pathway to sustain a steady-state flux through it. Such flux vectors are called Elementary Flux Modes (EFMs) [1]. Any positive steady-state flux vector may be expressed as a convex combination

$$(S3) \quad \mathbf{v} = \alpha_1 \mathbf{v}_1 + \alpha_2 \mathbf{v}_2$$

of these EFMs, with convex coefficients $\alpha_1$ and $\alpha_2$ that satisfy $0 \leq \alpha_1, \alpha_2 \leq 1$ and $\alpha_1 + \alpha_2 = 1$ [2].

#### S3 Derivation of the relation that determines the relative biomass composition

During balanced growth, intensive quantities such as concentrations of internal metabolites are in steady-state. The extensive quantities, such as the total biomass and volume, increase exponentially in time at a rate equal to the growth rate of the species. More specifically, the biomass of the species changes in time according to $B_A(t) = B_A^0 e^{\lambda_A t}$ and $B_B(t) = B_B^0 e^{\lambda_B t}$, where $B^0$ is the biomass of the species when the community attains a steady-state. Using these relations, we now reformulate the steady-state equation $\dot{P}_1 = 0$ from eq. (1) as

$$(S4) \quad \frac{B_A^0 e^{\lambda_A t}}{\rho_A} Y_{P_1/S_A^c} v_1^A = \frac{B_B^0 e^{\lambda_B t}}{\rho_B} v_U^B.$$

Since all quantities in this equation are either parameters or specific fluxes, which are intensive variables, the only time-dependent terms are the exponentially increasing functions. We use separation of variables to make sure that the time-dependent terms appear only on the left-hand side of the equation

$$(S5) \quad e^{(\lambda_A - \lambda_B)t} = \frac{B_B^0 \rho_A v_U^B}{B_A^0 \rho_B v_1^A} \frac{1}{Y_{P_1/S_A^c}}.$$

Since the right-hand side of the equation is independent of time, this equality only holds for all times if $\lambda_A = \lambda_B$, i.e., both community members grow equally fast. This is implied by assuming steady-state of $P_1$ and of the entire community.

However, explicitly using equality of the growth rates to model this community creates a problem when the producer uses exclusively mode 2 for growth, represented by $\mathbf{v}_2$ in eq. (S2). In that case, the producer does not excrete $P_1$, as the flux $v_1^A$ vanishes. Since $P_1$ is the carbon and energy source of the consumer, it cannot grow, so $\lambda_B = 0$. This breaks the growth rate equality.

To circumvent this modelling problem, we derive a different equation that implies growth rate equality when both species are present, but relaxes this condition when the producer uses mode 2 exclusively. Using the two steady-state relations for $X_B$ and ATP in the consumer from eq. (1), we rewrite the consumer's uptake flux as

$$(S6) \quad v_U^B = \frac{v_1^B + Y_{X_B^R/R_B} v_R^B}{Y_{X_B/P_1}} = \frac{\frac{m_B^B}{m_1^B} + Y_{X_B^R/R_B}}{Y_{X_B/P_1}} v_R^B.$$

Substituting this in eq. (S5) yields

$$(S7) \quad e^{(\lambda_A - \lambda_B)t} = \frac{B_B^0 \rho_A v_R^B \frac{m_B^B}{m_1^B} + Y_{X_B^R/R_B}}{B_A^0 \rho_B v_1^A Y_{P_1/S_A^c} Y_{X_B/P_1}}.$$

For this equation to hold at all times, it is again required that $\lambda_A = \lambda_B$. Furthermore, since the growth rate equals the protein synthesis rate per unit of protein ($\lambda = \frac{v_R}{e_{tot}}$), we use the growth rate equality to express the biomass synthesis rate of the consumer in that of the producer as

$$(S8) \quad v_R^B = \frac{e_{tot}^B}{e_{tot}^A} v_R^A.$$

Combining this with eq. (S7) gives

$$(S9) \quad 1 = \frac{B_B^0 \rho_A v_R^A e_{tot}^B \frac{m_B^B}{m_1^B} + Y_{X_B^R/R_B}}{B_A^0 \rho_B v_1^A e_{tot}^A Y_{P_1/S_A^c} Y_{X_B/P_1}},$$

that we rewrite as

$$(S10) \quad B_B^0 = \frac{v_1^A \rho_B e_{tot}^A Y_{P_1/S_A^c} Y_{X_B/P_1}}{v_R^A \rho_A e_{tot}^B \frac{m_B^B}{m_1^B} + Y_{X_B^R/R_B}} B_A^0.$$

Given a steady-state flux distribution (S3), this relationship determines the amount of steady-state biomass of the consumer ($B_B^0$) in terms of producer biomass ($B_A^0$). Therefore, we normalise the community biomass by setting $B_A^0 = 1$ gDW/L culture in the simulations.

The interpretation of eq. (S10) is as follows: when the producer invests less protein in mode 1, the flux ratio $\frac{v_1^A}{v_R^A}$ decreases. Since all other parameters on the right-hand side of the relation are constant, this leads to a lower relative biomass for the consumer ($B_B^0$). The intuition behind this is that the decrease in flux through mode 1 leads to a lower excretion rate of $P_1$, and thus a decrease in the uptake flux of $P_1$ to maintain steady-state. This results in a lower steady-state biomass of the consumer.

Similar behaviour occurs when we change one of the parameters. For instance, if the total enzyme concentration in the consumer ($e_{tot}^B$) increases for a constant flux ratio $\frac{v_1^A}{v_R^A}$, the consumer biomass decreases to satisfy eq. (S10). In this case the $P_1$ excretion flux remains constant while the uptake flux by the consumer increases per amount of biomass, due to its higher enzyme concentration. Attaining steady-state of $P_1$ thus requires less biomass of the consumer.

As was explained in Section 2.5, this relationship even holds when the producer uses mode 2 exclusively and does not excrete $P_1$ as the flux $v_1^A$ vanishes. In this case, the consumer cannot synthesise biomass, hence also $B_B^0$ vanishes. So, both sides of eq. (S10) are zero. This equation thus holds (and guarantees equal growth rates) when both species are present, but also when the consumer is not present. It is therefore a more general condition than the equality of the growth rates, which breaks down when only the producer is present.

### S4 Protein sectors and constraints in the model

The metabolic processes presented in Figure S1 are all associated with growth. As explained in Section 2.5, we model each of them by an enzymatic reaction $j$ that runs at rate $v_j = k_j e_j f_j(\mathbf{x})$, with corresponding enzyme concentration $e_j$. In addition to growth-associated processes, cells also spend a fraction of their enzymes on growth-unassociated processes such as maintenance, stress tolerance and preparation for future conditions [3–5]. The sum of the concentrations of growth-associated enzymes therefore does not add up to the total protein concentration ($e_{tot}$) in the cell. For both species modelled here this sum becomes, respectively,

$$(S11) \quad \begin{aligned} e_T^A + e_1^A + e_2^A + e_R^A &\leq e_{tot}^A \\ e_U^B + e_1^B + e_R^B &\leq e_{tot}^B. \end{aligned}$$

We assume that the total protein concentrations are equal for both species, to obtain a fair comparison, and that they are constant across conditions [6–8]. This constraint causes a trade-off between growth-associated and growth-unassociated processes [4, 9, 10]. Since this is not the focus of this work, we do not model the growth-unassociated processes and their enzyme concentrations explicitly.

For both the producer and the consumer we model similar catabolic and anabolic processes, but we model their assimilation of a carbon source in different ways. For the producer, the transport reaction ($T$) only represents import of the carbon source from the external environment to inside the cell. We assume that this import occurs through facilitated diffusion by a permease in the membrane. This is a passive process, meaning that it does not require any energy. As this transport reaction does not involve any chemical conversion, its equilibrium constant equals one. The Haldane relation [11] now directly couples the kinetic parameters of the permease $e_T^A$ in the membrane of the producer via

$$(S12) \quad \frac{k_T^+ K_{s_A}}{k_T^- K_s} = K_{eq}^T = 1.$$

This relation suggests a trade-off between the maximal rate and the substrate affinity of a permease [12]. To deal with this trade-off, cells typically have several different transporters for the same nutrient in their membrane. When substrate concentrations are high, the cell expresses transporters with low affinities, while high-affinity transporters are expressed in nutrient-scarce conditions [13, 14]. In our coarse-grained model, we only include one type of transporter in the producer, with a fixed affinity and catalytic rate constant. To account for the expression of different transporters that all occupy a fraction of the membrane space, we assume that the transporter concentration $e_T^A$ of the producer is constant across conditions. This assumption is also used in many other and more detailed models that study changes in metabolic strategies [9, 15–17].

For the consumer, the uptake process ($U$) contains both the import of the carbon source $P_1$ and its conversion to the intermediate $X_B$. The reason for this distinction is twofold. Firstly, as this uptake process contains a chemical conversion, eq. (S12) does not apply here. This allows us to adjust the affinity of the consumer for its carbon substrate independently of the other kinetic parameters related to uptake. Secondly, this uptake process describes more general transport processes than just facilitated diffusion, making the model more broadly applicable. This also means that the enzyme concentration $e_U^B$ for uptake is not constant in the consumer.

### S5 Solutions to the coupled optimisation problem

The derivations in Section 2.5 and the previous sections result in the optimisation problem for the community growth rate in eq. (5), that we restate here for completeness

$$(S13) \quad \max_{\mathbf{x}, \mathbf{e}, B_B^0} \left\{ \lambda_A = \frac{v_R^A}{e_{tot}^A} \left| \mathbf{N}\mathbf{v} = \mathbf{0}, e_T^A + e_1^A + e_2^A + e_R^A \leq e_{tot}^A, e_U^B + e_1^B + e_R^B \leq e_{tot}^B, \forall j : v_j = k_j e_j f_j(\mathbf{x}), e_j \geq 0, x_i \geq 0, \right. \right. \\ \left. \left. B_B^0 = \frac{v_1^A}{v_R^A} \frac{\rho_B}{\rho_A} \frac{e_{tot}^A}{e_{tot}^B} \frac{Y_{P_1/S_A^c} Y_{X_B/P_1}}{\frac{m_B^B}{m_1^B} + Y_{X_B^R/R_B}} B_A^0, \{S, P_2, P_B, B_A^0, e_T^A, e_{tot}^A, e_{tot}^B\} \text{ constant} \right\}.$$

A solution to this optimisation problem in terms of optimal metabolite and enzyme concentrations gives rise to an optimal flux vector $\mathbf{v}^{opt} = \mathbf{v}(\mathbf{x}^{opt}, \mathbf{e}^{opt})$ that is a convex combination of the EFMs of the metabolic network (cf. eq. (S3)). The optimal flux vector thus makes use of one or a combination of the catabolic pathways of the producer.

Earlier work by De Groot et al. [18] showed that the number of flux-carrying EFMs is bounded by the number of active flux-limiting constraints when a specific flux is optimised. Hence, to infer the metabolic strategy that maximises the growth rate in terms of the EFMs in eq. (S2), we need to examine the constraints in this model. In the previous section, we have identified these constraints in terms of enzyme concentrations. The producer is subject to two enzyme constraints: (i) its limited total enzyme concentration $e_{tot}^A$ and (ii) its constant transporter concentration $e_T^A$. The consumer also has a fixed total enzyme concentration $e_{tot}^B$, but since it has only one catabolic pathway that is only used when the producer excretes $P_1$, we can restrict our analysis to the two constraints for the producer.

Because we do not model growth-unassociated processes explicitly, maximisation of the growth rate results in a complete allocation of all enzymes to growth-associated processes in our model. In other words, the inequality for $e_{tot}^A$ in eq. (S11) will always become an equality. In reality, cells also spend a fraction of their enzymes on growth-unassociated processes, but we assume that this fraction is constant and negligible for the ecological phenomena described by our coarse-grained model. Including this would result in a quantitative change in enzyme allocation, but we are mainly interested in qualitative effects in this work. Spending all enzyme on growth-associated processes implies that two active flux-limiting constraints are active for the producer. Although one may expect that this also leads to a convex combination of the two flux-carrying EFMs as optimal flux vector, this does not need to be the case according to the extremum principle formulated by De Groot et al. [18]. Cells adjust their enzyme concentrations to tune fluxes, but these fluxes also depend on concentrations of metabolites through reaction kinetics. These extra degrees of freedom allow cells to tune fluxes within an EFM by changing the concentrations of enzymes, since the metabolite concentrations will change accordingly (up to some extent). This kinetic effect arising from metabolite variability can result in a single EFM as optimal flux vector, even though two constraints are active. Using less EFMs than the number of active constraints may thus be optimal in some cases [18]. In our model, the metabolite dependency appears in the saturation factors $f(\mathbf{x})$. In the following sections, we will see multiple instances in which the solution to eq. (S13) is a single EFM while two constraints are active.

The optimal flux vector may change when external conditions or physiological properties of community members change. In our framework, such altered conditions are represented by parameter perturbations. Varying a parameter independently thus describes a change in the community that affects its optimal metabolic strategy. This versatility allows us to capture different ecological phenomena with the same model. Throughout this work, we study the solution to the optimisation problem (S13) for different varying parameters and regimes.

### S6 Solution to the optimisation problem for a varying substrate concentration

Here, we first focus on varying the concentration of the carbon source of the producer. We set values for the model parameters as given in Table S1, that are chosen such that the cell can attain similar growth rates with both modes. Specifically, we set the ATP yield of catabolic pathway $2_A$ higher than that of pathway $1_A$, but give catabolic pathway $1_A$ a higher catalytic rate constant. In earlier work, we showed that these properties imply a higher biomass yield for mode 2, but a higher protein efficiency (flux per unit of protein)

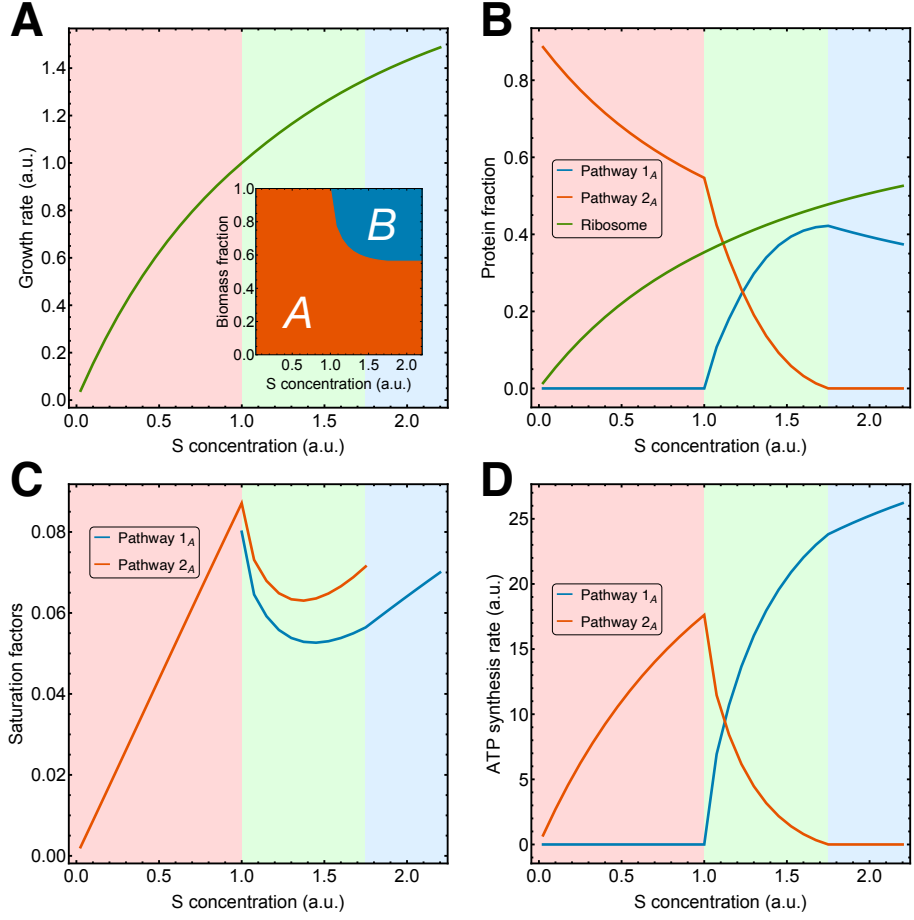

**FIGURE S2. Shift in the optimal catabolic pathway for increasing substrate concentration.** A gradual shift from mode 2 to mode 1 occurs as the substrate concentration $S$ increases beyond a critical value. This shift appears in **A.** the biomass composition of the community and **B.** the protein allocation in the producer. In the red region mode 2 is optimal, in the green region a combination of both modes is optimal and in the blue region mode 1 is optimal. When metabolite concentrations change such that **C.** the catabolic saturation factor of one mode increases, this results in **D.** a higher ATP synthesis rate with less enzyme. This occurs for mode 2 in the red region and for mode 1 in the blue region. In the green region, the enzyme limitation for mode 2 causes a shift in resource allocation toward mode 1 to increase the ATP synthesis rate further, since mode 1 has a higher protein efficiency. The growth rate and substrate concentration are normalised relative to the critical point where the shift initiates.

for mode 1 [9]. Experimental observations typically show that high-yield strategies are used at low substrate concentrations, while high-rate strategies are used at high substrate concentrations [19–21].

To verify that our model is consistent with these observations, we solve the optimisation problem numerically for a range of substrate concentrations. The resulting growth rate presented in Figure S2A increases hyperbolically with the substrate concentration, following the well-known Monod relation. For all substrate concentrations, the two enzyme constraints discussed in the previous section are active. Nevertheless, the protein fractions in Figure S2B show a change in the optimal metabolic strategy. In the red region at low substrate concentrations, where growth is mainly carbon-limited, the high-yield mode 2 reaches the highest growth rate [9]. Because the cell has spent all its enzymes on growth-associated processes, it faces a trade-off: to attain higher growth rates, the ribosomal protein fraction $e_R^A$ must increase at the expense of the catabolic fraction $e_2^A$. This poses a challenge for the cell, since a higher growth rate also requires a higher ATP synthesis rate by catabolism. However, the cell can accommodate this due to the kinetic effect of metabolite variability discussed in the previous section. Changes in metabolite concentrations increase the catabolic saturation factor $f_2$ and overcompensate for the loss of catabolic enzyme. This increases the rate of ATP synthesis by mode 2, as is depicted in the red region in Figures S2C and D.

Beyond a critical substrate concentration ($S_c$), the effect of metabolite variability on the saturation factor $f_2$ is not sufficient to deal with this trade-off using exclusively mode 2. In this green region, the optimal strategy gradually shifts from mode 2 to mode 1. Mode 1 requires more carbon but less protein to sustain the same ATP synthesis rate, thereby reaching higher growth rates when the carbon limitation relaxes at increasing substrate concentration. In this regime, the two active enzyme constraints do lead to a combination of two flux-carrying EFMs as optimal strategy. When all protein that was previously invested in mode 2 is completely used and replaced by mode 1, the optimal strategy is again a single EFM. In this blue region for $S > S_e$, the effect of metabolite variability on the saturation factor $f_1$ again allows the cell to achieve higher growth rates using mode 1 exclusively, despite the enzyme limitation. In summary, the behaviour of our model is indeed consistent with experimental observations for increasing substrate concentrations [19–21].

This analysis also illustrates that the number of active EFMs is determined by the substrate concentration. For $S < S_c$ and $S > S_e$, a single EFM maximises the growth rate, while for $S_c < S < S_e$, a convex combination of the EFMs does. We will exploit this model property to study different regimes that correspond to different ecological phenomena.

### S7 Solution to the optimisation problem for varying biomass yield of the consumer

In this section, we want to highlight one other parameter perturbation that results in a counterintuitive model prediction that can be tested experimentally. This is the biomass yield of the consumer, in our model presented by the parameter $Y_{X_B^R/R_B}^{-1}$. We analyse the effect of changing this yield solely for mode 1, since only this mode is affected by the consumer.

Increasing this parameter means that the consumer requires less of its carbon source $P_1$ to make one mole of biomass. One could now argue intuitively that this would lead to a rise in the external $P_1$ concentration, which would have a larger inhibitory effect on the producer, and therefore reduce the (community) growth rate. This intuitive reasoning is, however, incomplete. Because the biomass yield of the consumer increases (which means that the inverse yield $Y_{X_B^R/R_B}$ decreases), eq. (S10) requires that the consumer biomass $B_B^0$

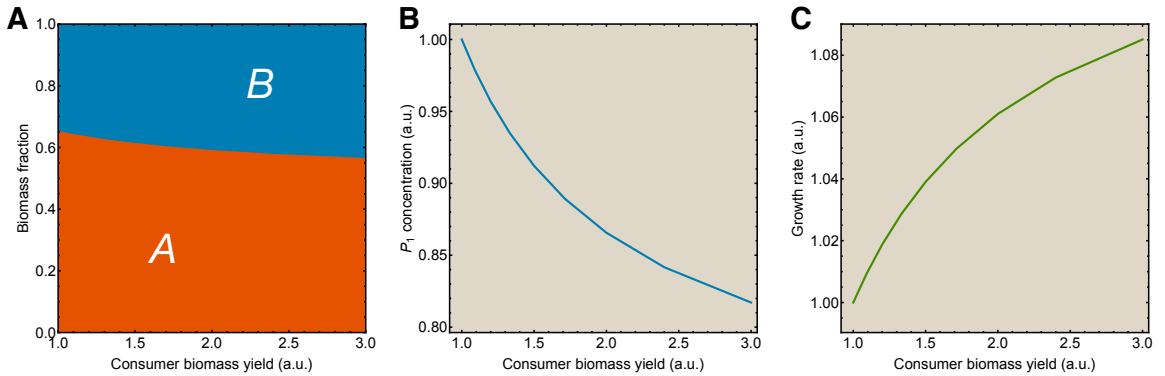

FIGURE S3. **Model results for varying biomass yield of the consumer.** As the biomass yield of the consumer increases, **A.** its relative biomass fraction increases, resulting in **B.** a lower $P_1$ concentration and **C.** a higher community growth rate.

increases as well. In biological terms, a higher biomass yield allows the consumer to synthesise more biomass per mole of $P_1$, increasing the relative biomass fraction of the consumer (Figure S3A). This increases the uptake rate of $P_1$, which leads to a drop in the steady-state concentration of $P_1$, as illustrated in Figure S3B. This results in a smaller inhibitory effect on the producer, which increases the growth rate for both community members, as is shown in Figure S3C.

### S8 Reduction of the optimisation problem for a passive consumer

In our model, the interaction between the producer and the consumer is mediated through the exchange of the byproduct $P_1$. This is an internal metabolite of the reaction network in Figure S1, so its (optimal) concentration is determined by the optimisation problem in eq. S13. A change in the properties of the consumer will thus only affect (the growth rate of) the producer if this results in a change in the optimal steady-state concentration of $P_1$. In this case, also the growth rate of the consumer is affected, since both species grow equally fast in a stable community.

It is insightful to formulate this reasoning the other way around. If an optimal $P_1$ concentration exists that is nonzero, this implies there exists a consumer phenotype that achieves the same growth rate as the producer at exactly this $P_1$ concentration. We may thus assume that such a consumer phenotype exists at any $P_1$ concentration that leads to a nonvanishing growth rate for both species. In other words, instead of optimising the growth rate of both species simultaneously (which is guaranteed via eq. (S10)), we may as well optimise the growth rate of the producer in isolation for a given $P_1$ concentration. We then assume that (a phenotype of) the consumer can achieve this growth rate at the given concentration of $P_1$. Since this approach does not require the optimisation of the growth rate of the consumer, we thus consider this species to be a *passive consumer* of the byproduct $P_1$. Hereby, we restrict our attention to the producer and its response to the consumer.

Like $S$ and $P_2$, also $P_1$ is now an external metabolite that is part of the (fixed) environment. We assume that the $P_1$ concentration that we fix explicitly is the steady-state concentration attained implicitly through diffusion and uptake by the consumer. So, we directly vary the $P_1$ concentration as an independent parameter. This reduces the reaction network in Figure S1 to the metabolic network of the producer alone, which is presented in Figure S4. This significantly simplifies the optimisation problem in eq. (S13) to

$$(S14) \quad \max_{e_A, S_A} \left\{ \lambda_A = \frac{v_R^A}{e_{tot}^A} \left| \mathbf{N}_A \mathbf{v}_A = \mathbf{0}, e_T^A + e_1^A + e_2^A + e_R^A \leq e_{tot}^A, \forall j : v_j = k_j e_j f_j(S_A), e_j \geq 0, S_A \geq 0, \right. \right. \\ \left. \left. \{S, P_1, P_2, e_T^A, e_{tot}^A\} \text{ constant} \right\}.$$

Here, the steady-state conditions $\mathbf{N}_A \mathbf{v}_A = \mathbf{0}$ describe the steady-states for $S_A$ and ATP in the producer.

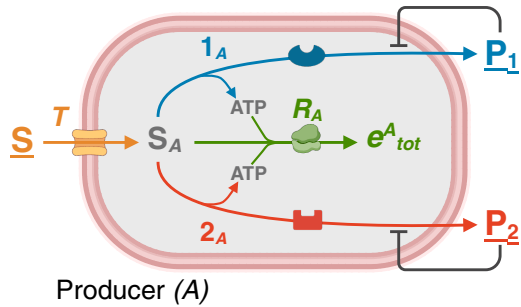

FIGURE S4. **Reduced metabolic network for the producer with a passive consumer.** Assuming that the consumer can attain the same growth rate as the producer for any positive concentration of $P_1$ reduces the metabolic network to that of the producer alone.

This reduction leads to the growth rate optimisation problem for a single species that is studied extensively in the literature [22]. The distinction here is, however, that we study this problem as function of the interaction between the community members via $P_1$. Similarly to the different ecological phenomena modelled by a variant of eq. (S13), a variant of this reduced model captures a simplified version of the phenomena discussed in this work. For each phenomenon, both approaches lead to qualitatively similar results.

### S9 Model variants to describe different positive ecological interactions

The general optimisation problem in eq. (S13) can be applied to study many different ecological scenarios that may occur in microbial communities. To analyse these scenarios systematically and couple them to observed phenomena, we use the following procedure. We start from a fixed microbial population as an initial condition. In most cases, this first population only contains the producer, so it is described by a subnetwork of the general metabolic network presented in Figure S1. For a fixed set of parameter values that describe the environment and the physiological properties of the species, we maximise the growth rate of this first population using the variant of the optimisation problem that corresponds to its subnetwork. The resulting growth rate serves as a reference point for comparison with a second population. This population is described by a different subnetwork that generally also contains a consumer, but using the same parameter values as the first population (at least for the parameters that appear in both subnetworks). Similarly as for the first population, we maximise the growth rate of the second population using the variant of the optimisation problem that corresponds to its subnetwork.

The key point is that we run the optimisation for the second population for varying values of one independent parameter. We are specifically interested in changes in the interaction between both species that affect their growth rate. We therefore pick the parameter that describes the affinity of the consumer for its carbon substrate $P_1$, which is a physiological property of the consumer. (In reality, the consumer will make use of different transporters with different affinities, as described in Section S4.) When this affinity increases, the consumer is better adapted to growing on the byproduct of the producer, as it can achieve a higher uptake rate at lower $P_1$ concentrations. This leads to an increased growth rate for both species, since a lower steady-state concentration of $P_1$ alleviates the inhibition on the producer. Increasing this affinity thus leads to an increased growth rate in the second population. For varying affinity, we then compare the growth rate of the second population with the (fixed) growth rate of the first population. When the growth rate of the second population exceeds that of the first population, this means that optimal community (composition) changes. If the second population also employs a different metabolic strategy than the first population, this is accompanied by a change in the optimal metabolic strategy.

Following this procedure, we use our framework to compare different populations. This systematic approach directly filters out the infeasible scenarios, which are situations in which the growth rate of the second population never exceeds that of the first population. Furthermore, our framework also excludes the possibility of two consuming species that both feed on the same carbon source. The consumer that achieves the highest growth rate will thrive. This is an example of competitive niche exclusion.

The remaining scenarios that we consider in this work correspond to the observed ecological phenomena discussed throughout the Results sections. Before discussing the specific model implementation for each phenomenon, we briefly discuss their main features. The parameters that remain fixed for all phenomena are given in Table S1. Since we study only qualitative effects in this work, these parameters are set to arbitrary values, while guaranteeing that the producer achieves similar growth rates for both modes.

The remaining parameters, which are the $S$ and $P_2$ concentrations, are set specifically for each phenomenon. For instance, for phenomena that involve a discrete switch in the optimal metabolic strategy we set $S < S_c$, as this ensures that the producer uses only one of the available metabolic modes to maximise its growth rate (cf. Section S6). Specifically for these phenomena that involve a change in the optimal metabolic strategy, we calibrate the reference growth rate attained by mode 2 by adjusting the concentration of its byproduct $P_2$ in the external environment. For each phenomenon, we have summarised the specific choices for the first population, the second population and parameter values in Table S2. These are used in the Mathematica implementations of the model, which are provided in the source code on Zenodo. Other details are discussed in the following sections.

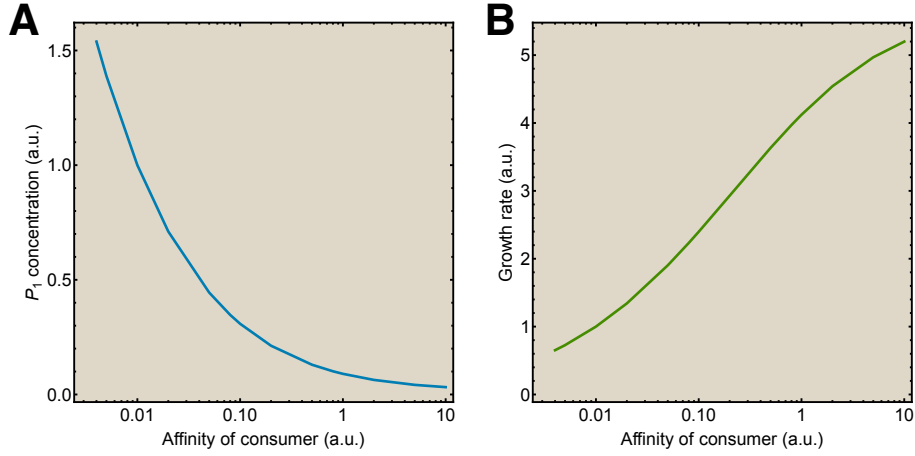

**FIGURE S5. Increasing the affinity of the consumer gives rise to enhanced mutualistic interactions.** As the affinity of the consumer for its carbon substrate $P_1$ increases, this **A.** decreases the steady-state concentration of $P_1$, which **B.** increases the community growth rate.

### S10 Model and implementation for mutualism and obligate syntrophy

The subnetworks used to describe the populations that are associated with mutualism are shown in Figures 3A and B. Because mode 2 of the producer is not present in these metabolic networks, we use a variant of the optimisation problem (S13) that omits this mode. Since both populations use the same metabolic mode, we set an arbitrary reference point that represents the first population containing only the producer, to which we normalise the $P_1$ concentration and the growth rate. For the second community containing both the consumer and the producer, we increase the affinity of the consumer for its carbon substrate, which results in a lower $P_1$ concentration and thus a higher community growth rate. This is depicted in Figure S5.

In fact, as mentioned in Section 2.2, the community with the consumer will always lower the steady-state $P_1$ concentration with respect to the population without the consumer, since it includes consumption of $P_1$. The growth rate of the community with the consumer will therefore always exceed that of community without the consumer. (Provided that the affinity of the consumer for $P_1$ is sufficiently high so that it can grow at this steady-state concentration). Combining Figure S5A and Figure S5B as function of the affinity then results in Figure 3C.

To model obligate syntrophy as discussed in Section 2.3, we use the same metabolic networks and optimisation problem as for mutualism, but study these close to thermodynamic equilibrium. Since we only include concentrations of catabolic metabolites in our model, this regime is defined by the distance from thermodynamic equilibrium for the catabolic pathway. As the $P_1$ concentration increases, e.g. through accumulation, this pathway gets closer to its equilibrium.

In practice, this means that we evaluate the optimisation problem at low values for the affinity of the consumer (compared to the values in Figure S5). Lowering the affinity results in a higher steady-state concentration for $P_1$, bringing it closer to its equilibrium concentration, as shown in Figure S6A. This decreases the catabolic flux in the producer, which can only be compensated by increasing the catabolic enzyme fraction. Due to the trade-off between catabolic and anabolic enzyme allocation, this also leads to a drop in growth rate, presented in Figure S6B. For very low affinity, there is hardly any uptake by the consumer. This approximates a situation in which there is no consumption, as in the community without the consumer, and thus $P_1$ accumulates in the medium until it reaches its equilibrium concentration. At this stage, the growth rate drops to zero. For any community where there is a consumer with nonzero affinity, additional consumption of $P_1$ results in a lower steady-state concentration and thus a nonzero growth rate.

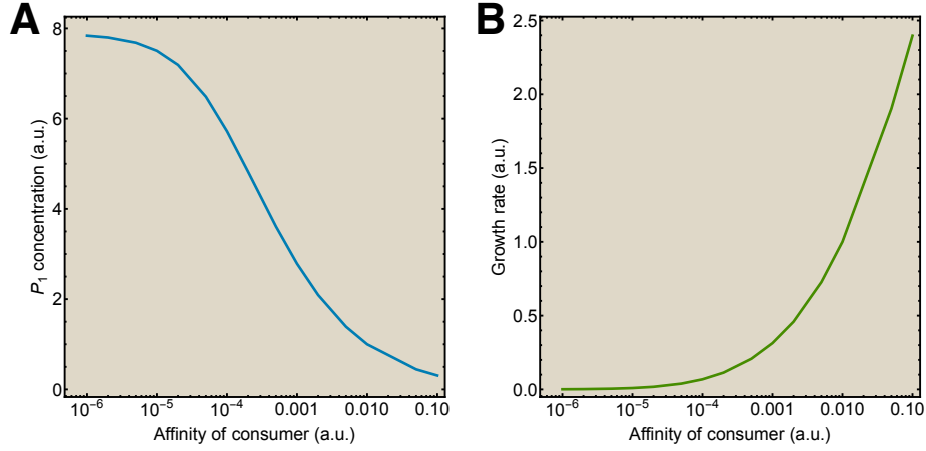

**FIGURE S6. When $P_1$ accumulates, the community attains thermodynamic equilibrium.** As the affinity of the consumer for its carbon substrate $P_1$ decreases, this **A.** increases the steady-state concentration of $P_1$, which **B.** decreases the community growth rate. For very low affinity, this simulates a scenario without consumption where the producer attains thermodynamic equilibrium due to accumulation of $P_1$.

### S11 Model and implementation for facultative syntrophy

For facultative syntrophy, the corresponding subnetworks that describe the different populations are shown in Figures 5A and B. Since both catabolic pathways of the producer are present in these metabolic networks, we use the complete optimisation problem (S13) to model this phenomenon. We set the external concentration of $P_2$ to a value such that the producer in isolation achieves a higher growth rate using mode 2 when the steady-state concentration of $P_1$ is high. This means that a community that uses mode 1 and contains consumers with a low affinity will grow slower than the producer on its own using mode 2. When the affinity of the consumer exceeds a threshold value, the steady-state $P_1$ concentration has dropped sufficiently for the producer to achieve a higher growth rate using mode 1. Since we set $S < S_c$, the presence of the consumer causes a discrete switch in the optimal metabolic strategy from mode 2 to mode 1 at this critical cooperation point, as indicated in Figures 5C and D.

Using the model reduction detailed in Section S8 we also simulate facultative syntrophy by explicitly varying the $P_1$ concentration. Similarly to varying the affinity, this leads to a switch in the optimal strategy from mode 2 to mode 1, as depicted in Figure S7. This confirms that the reduced model indeed leads to qualitatively similar results as the complete model for facultative syntrophy. However, the regimes in Figure S7 are reversed with respect to Figure 5C, since a higher affinity results in a lower $P_1$ concentration. Also it should be noted that in the red region, the $P_1$ concentration is a ghost parameter, since this byproduct is not synthesised when mode 2 is used exclusively.

This fact is also important when we consider the dynamics during the invasion by the community consisting of both producers and consumers. In this work, we do not consider these dynamics explicitly. When the producer appears in isolation, it will only use mode 2 and excrete byproduct $P_2$. In this environment, phenotypes of the producer that use mode 1 and excrete $P_1$ will be outcompeted, as they grow slower. Nevertheless, there will always be a few producer cells that use mode 1 due to genetic mutations and environmental fluctuations. For these cells, there will be a (local) quasi steady-state $P_1$ concentration around them. If these cells are close to some consumer cells that consume $P_1$ and decrease its external concentration, this leads to a higher growth rate of the producer phenotypes using mode 1. When the growth rate of these cells exceeds that of the producer cells using mode 2, the mode 1-phenotype together with the consumers will grow in relative abundance. This is how the community containing both the producer and the consumer can invade the population with only producers.

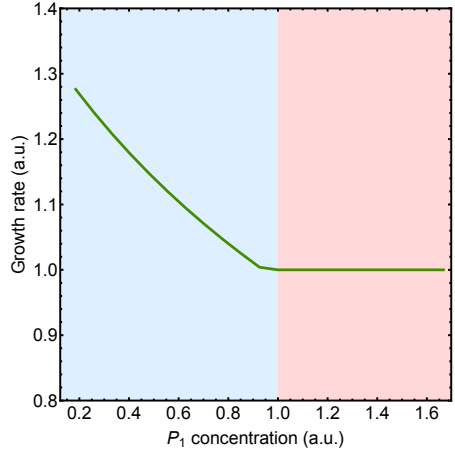

FIGURE S7. **Switch in the optimal catabolic pathway also arises in the reduced model with a passive consumer.** As the concentration of $P_1$ decreases below a critical concentration, the producer switches its optimal strategy from catabolic mode 2 (red region) to mode 1 (blue region). This results in a switch in the biomass composition of the community and an increasing community growth rate (normalised relative to the population containing only the producer).

### S12 Model and implementation for division of labour

The reaction networks used to model division of labour (DOL), which are shown in Figures 6A and B, differ slightly from the general metabolic network in Figure S1. This is because we compare a complete degradation of the substrate $S$ to $P_2$ by the producer to a DOL strategy that splits the degradation between the producer and the consumer. Since both these catabolic pathways describe the same chemical conversions, they consist of the same number of reactions, which we take to be three in this model. We include this in the model by extending mode 2 with two additional reactions. For both modes, reaction $1_A$ occurs in the producer, which also describes excretion in the case of the DOL strategy, just like in the other models. The two other catabolic reactions occur either in the producer for mode 2 or in the consumer for mode 1. For these reactions the same chemical conversions occur, but under different conditions and physiological properties, which means that the kinetics of these reactions are also different in both species. This allows us to freely choose the kinetic parameters for these two catabolic reactions in both species. The implementation of the optimisation problem for these larger metabolic networks and the corresponding extra parameters and their values can be found in the source code on Zenodo.

Increasing the affinity of the consumer again results in a switch in the optimal metabolic strategy after a threshold value, as shown in Figure S8. After this threshold, the growth rate achieved using the DOL strategy exceeds the growth rate achieved using the complete degradation by the producer. Figure S8 is qualitatively identical to Figures 5C and D for facultative syntrophy. This demonstrates that DOL may be interpreted as a special case of facultative syntrophy. Also, this analysis shows that DOL may arise solely from differences between both species in kinetic parameters of reactions.

In Figures 6C and D we showed that DOL also becomes the favourable strategy at higher substrate concentrations in our model, which is consistent with experimental observations and other theoretical predictions [19, 20, 23]. When the substrate concentration exceeds a critical value, the optimal strategy changes gradually from complete degradation to the DOL strategy. The reason that this shows a gradual shift instead of an abrupt switch is the same as explained in Section S6. Indeed, Figures 6C and D are qualitatively identical to Figures S2A and B. Because we use a different metabolic network to model DOL, the value for the critical substrate concentration $S_c$ differs in both simulations.

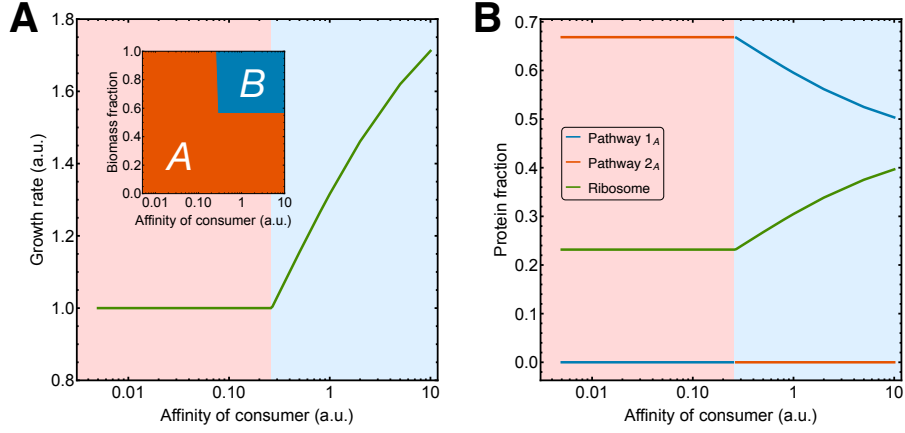

**FIGURE S8. Switch from complete degradation to DOL for increasing affinity of the consumer.** A discrete switch from complete degradation by the producer (red region) to DOL by both species (blue region) occurs as the affinity of the consumer for $P_1$ increases beyond a critical value. This results in **A**, an increase in the growth rate (normalised relative to the population containing only the producer) and a switch in the biomass composition of the community and **B**, a switch in the protein allocation in the producer.

#### S13 Model and implementation for syntrophic interactions with multiple consumers

In the first part of Section 2.6 we model two communities in which a consumer of the byproduct of the producer is present. Each community uses one of the two available modes. These are presented in Figures 7A and B. We model the additional consumer ($C$) of $P_2$ by a similar metabolic network as the consumer ($B$) of $P_1$. This also requires an extension of the optimisation problem (S13) such that the steady-state equations for $P_2$ and $X_C$, the constraint and the total protein content and another biomass relation between the species are also taken into account. Since these additional equations and constraints are similar to consumer $B$, they are not stated explicitly here. The properties of species $C$ are picked such that the community consisting of $A$ and $C$ attains a similar growth rate as that consisting of $A$ and $B$. The exact model with its parameters and the implementation can be found in the source code on Zenodo.

Since the steady-state $P_2$ concentration in this community arises from both diffusion and consumption by species $C$, it is lower compared to its (fixed) steady-state concentration in the model for facultative syntrophy, where only diffusion has an effect. To make this difference apparent, we use the growth rate of the population with only the producer from Figure 5C as a reference point in Figure 7C. Indeed, the growth rate of the community in which consumer $C$ is present is higher due to the lower $P_2$ concentration. As a result, the condition for invasion by the community with species $B$ becomes more restrictive, meaning that it can only grow faster at a higher affinity. This is demonstrated by the increase in the threshold value of the affinity, which determines the onset of the switch in the optimal strategy.

#### S14 Model and implementation for resource allocation over mixed strategies

In the second part of Section 2.6 we study a scenario in which the optimal metabolic strategy of the producer in isolation is a convex combination (S3) of both modes. To model this situation, we use the constraint analysis from Sections S5 and S6 to set the fixed substrate concentration to a value that exceeds the critical substrate concentration $S_c$. In this regime, the two active growth-limiting constraints on the producer also result in an optimal metabolic strategies that combines both modes.

Given this population as initial condition, we introduce the consumer of $P_1$ in the second community. Increasing the affinity of the consumer then leads to a decrease in the concentration of $P_1$, and thus an increase in the relative protein investment in mode 1 by the producer, as explained and shown in Section 2.6.

Although the increased protein investment in mode 1 results in a higher excretion rate of $P_1$, the increased relative abundance of the consumer (cf. the small panel in Figure 8C) compensates for this, leading to an overall decrease in the steady-state concentration of $P_1$. This increases the community growth rate due to the presence of the consumer. The consumer of $P_1$ can thus always invade in this situation. A consumer of $P_2$ that decreases the steady-state concentration of this byproduct would also be able to invade the population with only producers in a similar way.

In mathematical terms, increasing the consumers affinity for $P_1$ increases the convex coefficient of mode 1, which results in a drop in the convex coefficient of mode 2. For a significant increase in the consumers affinity, the convex coefficient of mode 1 equals 1, meaning that the optimal metabolic strategy is given by mode 1 exclusively. We also observed this for high substrate concentrations ($S > S_e$) in Figure 6D and Figure S2B. For very low affinity, the optimal strategy is given by mode 2 exclusively. In both these regimes, the two active growth-limiting constraints do not result in two active metabolic strategies, which follows from the metabolite variability effect in this model (cf. Section S5).

| Parameter symbol | Parameter description | Value |
| --- | --- | --- |
| $B_0^A$ | Fixed biomass of the producer | 1 gDW/L culture |
| $e_{tot}^A$ | Total protein concentration in the producer | 1 mmol protein/L cell |
| $e_{tot}^B$ | Total protein concentration in the consumer | 1 mmol protein/L cell |
| $e_T^A$ | Fixed transporter protein concentration in the producer | 0.1 mmol protein/L cells |
| $f_R^A$ | Saturation factor of the ribosome in the producer | $0.8 \text{ h}^{-1}$ |
| $f_R^B$ | Saturation factor of the ribosome in the consumer | $0.8 \text{ h}^{-1}$ |
| $k_T^A$ | Catalytic rate constant of carbon transport in the producer | $1000 \text{ h}^{-1}$ |
| $k_1^A$ | Catalytic rate constant of catabolic pathway $1_A$ in the producer | $1000 \text{ h}^{-1}$ |
| $k_2^A$ | Catalytic rate constant of catabolic pathway $2_A$ in the producer | $185 \text{ h}^{-1}$ |
| $k_R^A$ | Catalytic rate constant of the ribosome in the producer | $0.089 \text{ h}^{-1}$ |
| $k_U^B$ | Catalytic rate constant of carbon uptake in the consumer | $1000 \text{ h}^{-1}$ |
| $k_1^B$ | Catalytic rate constant of the catabolic pathway in the consumer | $500 \text{ h}^{-1}$ |
| $k_R^B$ | Catalytic rate constant of the ribosome in the consumer | $0.089 \text{ h}^{-1}$ |
| $K_{eq}$ | Equilibrium constant for catabolic reactions | 10 (100 for varying $S$) |
| $K_{eq}^T$ | Equilibrium constant for transport by the producer | 1 |
| $K_s$ | Michaelis constant for $S$ | 1 mM |
| $K_{s_A}$ | Michaelis constant for $S_A$ | 2 mM |
| $K_{p_1}^A$ | Michaelis constant for $P_1$ in the producer | 0.01 mM (1 for varying $S$) |
| $K_{p_2}$ | Michaelis constant for $P_2$ | 1 mM |
| $K_{x_B}$ | Michaelis constant for $X_B$ | 1 mM |
| $K_{p_B}$ | Michaelis constant for $P_B$ | 2 mM |
| $m_1^A$ | ATP yield of catabolic pathway $1_A$ in the producer | 1 mmol ATP/mmol $S$ |
| $m_2^A$ | ATP yield of catabolic pathway $2_A$ in the producer | 2 mmol ATP/mmol $S$ |
| $m_R^A$ | ATP requirement per mmol protein in the producer | 700 mmol ATP/mmol protein |
| $m_1^B$ | ATP yield of catabolic pathway $1_B$ in the consumer | 1 mmol ATP/mmol $X_B$ |
| $m_R^B$ | ATP requirement per mmol protein in the consumer | 700 mmol ATP/mmol protein |
| $P_B$ | Fixed concentration of byproduct $P_B$ | 0.02 mM (0.2 for varying $S$) |
| $Y_{S_A/R_A}$ | Carbon requirement per mmol protein for the producer | 200 mmol $S$/mmol protein |
| $Y_{X_B/R_B}$ | Carbon requirement per mmol protein for the consumer | 200 mmol $X_B$/mmol protein |
| $Y_{P_1/S_A^c}$ | $P_1$ yield per $S$ | 1 mmol $P_1$/mmol $S$ |
| $Y_{X_B/P_1}$ | $X_B$ yield per $P_1$ | 1 mmol $X_B$/mmol $P_1$ |

TABLE S1. **Fixed parameter values used in model variants for different cooperative ecological phenomena.** Parameter values that are used in the Mathematica implementations of the models, which are provided in the source code on Zenodo. For the models that describe DOL and multiple consumers, we use a larger metabolic network, as explained in Sections S12 and S13. These models contain additional parameters, that are specified in the source code.

| Phenomenon | First population<br>(metabolic mode) | Second population<br>(metabolic mode) | Specific fixed pa-<br>rameters |
| --- | --- | --- | --- |
| 2.2 Mutualism<sup>a</sup> | $A$ (1) | $A + B$ (1) | $S = 0.3 \text{ mM} (< S_c)$ |
| 2.3 Obligate<br>syntrophy<sup>a</sup> | - | $A + B$ (1) | $S = 0.3 \text{ mM} (< S_c)$ |
| 2.3 Facultative<br>syntrophy | $A$ (2) | $A + B$ (1) | $S = 0.3 \text{ mM} (< S_c)$,<br>$P_2 = 0.5 \text{ mM}$ |
| 2.4 Division of<br>labour (DOL)<sup>b</sup> | $A$ (2) | $A + B$ (1) | $S = 0.8 \text{ mM} (< S_c)$,<br>$P_2 = 0.01 \text{ mM}$ |
| 2.6 Syntrophy<br>(two consumers) | $A + C$ (2) | $A + B$ (1) | $S = 0.35 \text{ mM} (< S_c)$ |
| 2.6 Mixed strate-<br>gies | $A$ (1+2) | $A + B$ (1+2) | $S = 0.5 \text{ mM} (> S_c)$,<br>$P_2 = 0.01 \text{ mM}$ |

TABLE S2. **Specific model variants and parameter values for different cooperative ecological phenomena.** <sup>a</sup>: For mutualism and obligate syntrophy, we focus exclusively on mode 1, so mode 2 and its byproduct $P_2$ do not appear here. <sup>b</sup>: For DOL, we use a larger metabolic network with multiple catabolic reactions. This is explained in full detail in Section S12.
